# Mathematical framework for place coding in the auditory system

**DOI:** 10.1101/2020.05.20.107417

**Authors:** Alex D. Reyes

## Abstract

In the auditory system, tonotopy is postulated to be the substrate for a place code, where sound frequency is encoded by the location of the neurons that fire during the stimulus. Though conceptually simple, the computations that allow for the representation of intensity and complex sounds are poorly understood. Here, a mathematical framework is developed in order to define clearly the conditions that support a place code. To accommodate both frequency and intensity information, the neural network is described as a topological space with elements that represent individual neurons and clusters of neurons. A bijective mapping is then constructed from acoustic space to neural space so that frequency and intensity are encoded, respectively, by the location and size of the clusters. Algebraic operations -addition and multiplication- are derived to elucidate the rules for representing, assembling, and modulating multi-frequency sound in networks. The predicted outcomes of these operations are consistent with network simulations as well as with electrophysiological and psychophysical data. The analyses show how both frequency and intensity can be encoded with a purely place code, without the need for rate or temporal coding schemes. The algebraic operations are used to describe loudness summation and suggest a mechanism for the critical band. The mathematical approach complements experimental and computational approaches and provides a foundation for interpreting data and constructing models.

**Author Summary:** One way of encoding sensory information in the brain is with a so-called place code. In the auditory system, tones of increasing frequencies activate sets of neurons at progressively different locations along an axis. The goal of this study is to elucidate the mathematical principles for representing tone frequency and intensity in neural networks. The rigorous, formal process ensures that the conditions for a place code and the associated computations are defined precisely. This mathematical approach offers new insights into experimental data and a framework for constructing network models.

## Introduction

Many sensory systems are organized topographically so that adjacent neurons have small differences in the receptive fields. The result is that minute changes in the sensory features causes an incremental shift in the spatial distribution of active neurons. This is has led to the notion of a place code where the location of the active neurons provides information about sensory attributes. In the auditory system, the substrate for a place code is tonotopy, where the preferred frequency of each neuron varies systematically along one axis [1]. Tonotopy originates in the cochlea [2, 3] and is inherited by progressively higher order structures along the auditory pathway [4]. The importance of a place code [5] is underscored by the fact that cochlear implants, arguably the most successful brain-machine interface, enable deaf patients to discriminate tone pitch simply by delivering brief electrical pulses at points of the cochlea corresponding to specific frequencies [6].

Although frequency and intensity may be encoded in several ways [7], there are regimes where place-coding seems advantageous. Humans are able to discriminate small differences in frequencies and intensities even for stimuli as brief as 5-10 ms [8, 9, 10, 11, 12]. Therefore, the major computations have already taken place within a few milliseconds. This is of some significance because in this short time interval, neurons can fire only bursts of 1-2 action potentials [13, 14], indicating that neurons essentially function as binary units. Therefore, it seems likely that neither frequency nor intensity can be encoded via the firing rate of individual cells since the dynamic range would be severely limited. Similarly, coding schemes based on temporal or ‘volley’ schemes is difficult to implement at the level of cortex because neurons can phase-lock only to low frequency sounds [15, 16, 17].

There are, however, several challenges with implementing a place coding scheme. First, the optimal architecture for representing frequency is not well-defined. Possible functional units include individual neurons, cortical columns [18, 19], or overlapping neuron clusters [20]. The dimension of each unit ultimately determines the range and resolution at which frequencies and intensities that can be represented and discriminated. Second, how both frequency and intensity can be encoded with a place code is unclear, particularly for brief stimuli when cells function mostly as binary units. Other variables such as firing rate or spike timing are commonly incorporated into models [21]. Third, the rules for combining multiple stimuli is lacking. Physiological sounds are composed of pure tones with differing frequencies and intensities, resulting in potentially complex spatial activity patterns in networks. Finally, the role of inhibition in a place coding scheme has not been established.

Here, a mathematical model is developed in order to gain insights into: 1) the functional organization of the auditory system that supports a place coding scheme for frequency and intensity: and 2) the computations that can be performed in networks. The approach is to use principles from set theory and topology to construct the acoustic and neural spaces, find a mapping between the spaces, and then develop the algebraic operations. The predictions of the math model are then tested with simulations.

## Results

The mathematical model is subject to the following biological constraints. First, the neural network inherits the tonotopic organization of the cochlea [2, 3] so that the preferred frequency of neurons changes systematically with location along one axis. Second, a pure tone activates a population of neurons within a confined area [22], with the location of the area varying with the tone frequency. Third, the area grows with sound intensity [22], paralleling the increase in the response area of the basilar membrane [23, 24]. The model is broadly related to the “spread-of-excitation” class of models [7, 25]

The basic computations that can be performed using a place code are shown schematically in a 2-dimensional network of neurons (Fig. 1). In response to a pure tone stimulus, a synaptic field from a presynaptic population of neurons is generated within an enclosed area of the network (a, left panel, cyan disk), causing a population of cells to fire (filled circles). A tone with a higher frequency and intensity activates a larger area at a different location (right panel). A sound composed of the two pure tones activates both regions simultaneously; the regions may overlap if the difference in frequencies is small (Fig. 1b). Finally, excitatory synaptic fields and the activated neuron clusters are modulated by inhibition (Fig. 1c). The mathematical basis for these computations is developed below. For clarity, only the main results are shown (details in Supporting Text S1).

**Figure 1:**
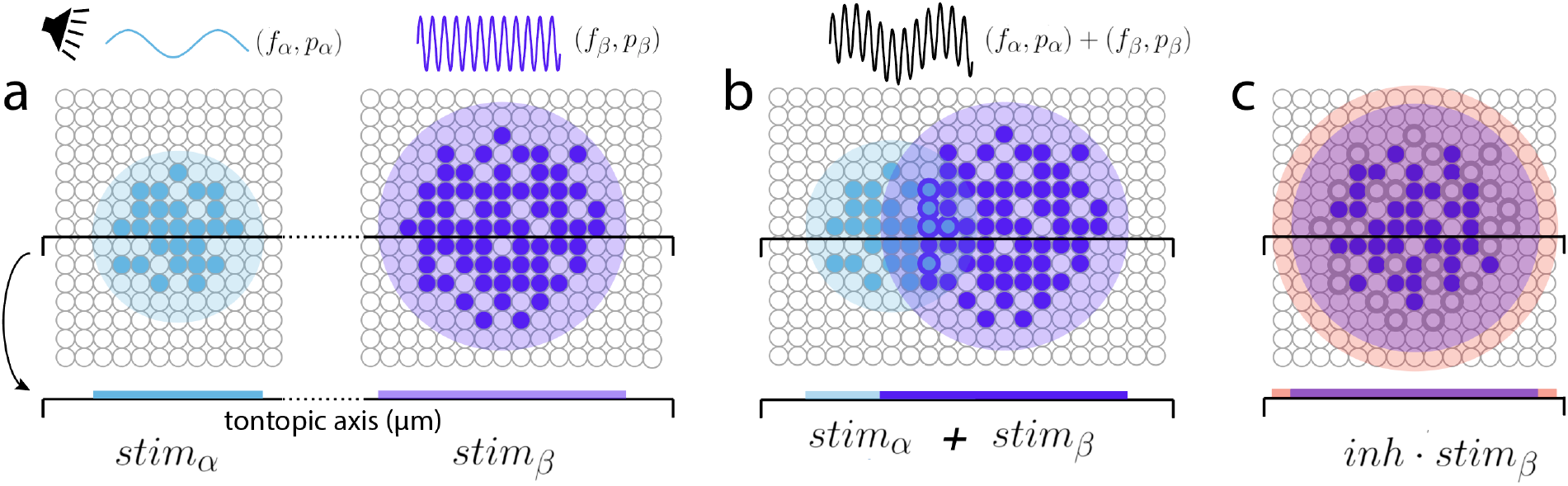
Computations with a place code. ***a, left***, hypothetical neural representation of a low frequency (*f_α_*), small amplitude (*p_α_*) pure tone stimulus in a two-dimensional neural network. A stimulus-evoked synaptic field covers a circular area (cyan disk) and causes a subset of cells to fire (filled circles). Projection of the synaptic field onto the tonotopic axis (cyan bar) gives the location and size of the activated area. ***right***, synaptic field generated by a tone with higher frequency (*f_β_*) and sound pressure (*p_β_*). ***b***, synaptic field generated by a sound composed of the two tones. ***c***, modulation by inhibition (red).

### Neural Space

Although the brain has three spatial dimensions, only one dimension-that corresponding to the tonotopic axis-is relevant for a place code. Thus, the circular synaptic field in Fig. 1 is projected onto the tonotopic axis (bars). For simplicity, the neural space is defined as a single row of neurons (Fig. 2a). In the presence of sound, afferents from an external source generates a synaptic field that covers a contiguous subset of neurons. In the following, a mathematical description of the neural space will be developed that accommodates the neural elements and synaptic fields.

**Figure 2:**
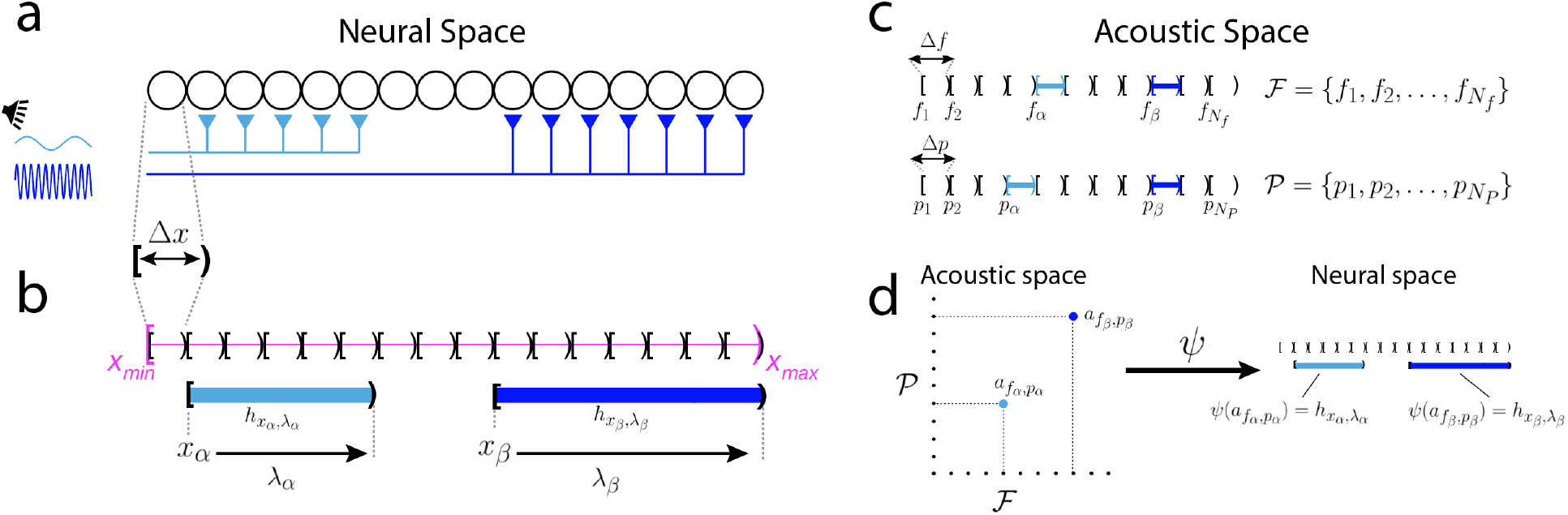
Mathematical representation of neural and acoustic Spaces. ***a***, neurons on the tonotopic axis are positioned next to each other with no space in between. Pure tone stimuli activate afferents onto a subset of neurons (cyan, blue). ***b***, mathematical representation of neural space. The tonotopic axis is a half-open interval (magenta) partitioned into smaller intervals that represent the space (Δ*x*) taken up by neurons. The synaptic fields 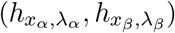 are also halfopen intervals. ***c***, The acoustic space has two dimensions, with frequency as one axis and pressure as the other. The frequency and pressure spaces are partitioned into half-open intervals of length Δ*f*, Δ*p*, respectively. ***d***, Mapping two tones in the acoustic space to intervals in neural space via a function *ψ*.

The neural space is defined as an interval, bounded by minimum and maximum values *x_min_* and *x_max_* (Fig. 2b, magenta). To incorporate neurons, the neural space is partitioned into *N_cell_* sections. Both the neural space and the cells are ‘half-open’ intervals, which are closed (“[”) on one end and open on the other (“)”). This is convenient mathematically because neurons can be packed next to each other with no space in between, allowing for a formal definition of a partition (see Supporting Text S1). Each neuron can therefore be expressed as non-overlapping subintervals of the form [*x, x* + Δ*x*), where Δ*x* is the width of each neuron. Note that each neuron can be uniquely identified by the point at the closed end, which also gives its location along the tonotopic axis. The set containing these points is:

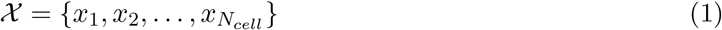

A synaptic field spans an integral number (*n*_λ_) of neurons and is also defined as a half-open interval with length λ = *n*_λ_Δ*x*. The length λ ranges from Δ*x* (1 cell) to a maximum λ_*max*_ = *n*_λ,*max*_Δ*x*. Each synaptic interval, designated as *h*_*x*,λ_ = [*x, x* + λ), is uniquely identified by the location of the cell at the closed end (*x_α_, x_β_* in Fig. 2b) and by its length (λ_*α*_, λ_*β*_). The set of starting points 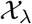 and the set of achievable lengths Λ are given by:

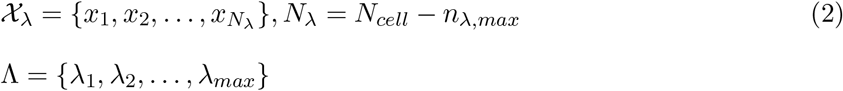

As will be shown below, 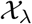 will contain information about the frequency while Λ will contain information about the sound pressure.

The set of all synaptic intervals in neural space is a basis for topology on the neural space, which will be used to define algebraic operations and construct mapping from acoustic space (see below, Supporting Text S1).

### Representing sound in neural space

The elements of acoustic space 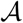 are pure tones, each of which is characterized by a sine wave of a given frequency (*f*) and amplitude corresponding to sound pressure (*p*). Theoretically, frequencies and pressures are unbounded and can take an uncountable number of values but under physiological conditions, the audible range is likely bounded by minimum and maximum values and is finite.

The acoustic space has two dimensions with frequency as one axis and sound pressure as the other. As was done for neural space, the frequency and pressure axes are defined as a half-open intervals and divided into non-overlapping subintervals expressed as [*f, f* + Δ*f*) and [*p, p* + Δ*p*), respectively (Fig. 2c). Physiologically, Δ*f* and Δ*p* is loosely related the minimum frequency and pressure difference that can be discriminated (frequency and intensity difference limens, see Discussion) [8, 9, 10, 11, 12]). For example, two tones with frequencies *f_α_, f_β_* that are in the interval [*f*_1_, *f*_1_ + Δ*f*) will both be ‘assigned’ to *f*_1_ (see Supporting Text S1). Therefore, the sets of audible frequencies and pressures are given by the first points of each interval:

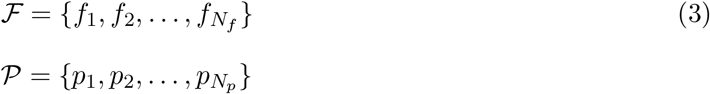

The number of audible frequencies and pressures are limited by the number of synaptic intervals that fit into the tonotopic axis (*N*_λ_ in eq. 3 and the maximum number of cells that fit into a single synaptic interval (*n*_λ, *max*_), respectively.

Features of the acoustic space are represented in neural space via a mapping *ψ* (Fig. 2d; see Supporting Text S1 for formal treatment). A pure tone 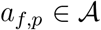 is mapped to an interval *h*_*x*,λ_ by first mapping the components *f* to *x* and *p* to λ via *ψ_f_*(*f*) = *x* and *ψ_p_*(*p*) = λ.

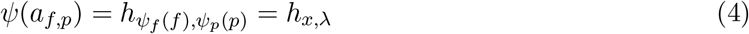

By adjusting the number of elements in frequency and pressure spaces to match the number of synaptic intervals (*N*_λ_) and maximum number of cells within a synaptic interval (*n*_λ,max_), respectively, the mapping becomes bijective (one-to-one, onto).

A mapping from acoustic space to intervals that are inhibitory can be similarly defined. The mapping is complicated by the fact that the number and receptive field properties of excitatory (***E***) and inhibitory (***I***) cells differ. The mapping is under ongoing investigation but for purposes of the present analyses, the mapping is taken to be identical to that for ***E***.

### Algebraic operations with synaptic intervals

Having formally described the mathematical structure of neural space, it is now possible to define the algebraic operations-addition and multiplication-for combining and modulating synaptic intervals (Fig. 1b,c). An important condition that must be satisfied is that operations be ‘closed’; that is, the result obtained with addition or multiplication is an interval (or intervals) that remains in neural space. To simplify notation, the intervals will henceforth be identified with a single subscript or superscript.

To satisfy closure, ‘addition’ of synaptic intervals is defined as their union. Let *h_α_, h_β_* be two synaptic intervals. Then,

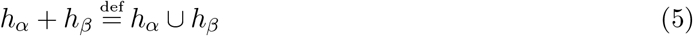

The addition operation yields two possible results (see Supporting Text S1 for details). If the two synaptic intervals do not overlap (*h_β_*, *h_γ_* in Fig. 3a*i*), the union yields a set with the same two intervals (Fig. 3a*ii*, bottom). If the two intervals overlap (*h_α_* and *h_β_*, *h_α_* and *h_γ_*), the result is a single interval whose length depends on the amount of overlap (Fig. 3a*ii*, top two traces). Note that summation is sublinear: e.g. |*h_α_* + *h_γ_*| ≤ |*h_α_*| + |*h_γ_*|. Moreover, if the starting points are the same (*h_α_* and *h_β_*), the length is equal to that of longer summand (|*h_α_* + *h_β_*| = |*h_α_*|, |*h_α_*| ≥ |*h_β_*|).

**Figure 3:**
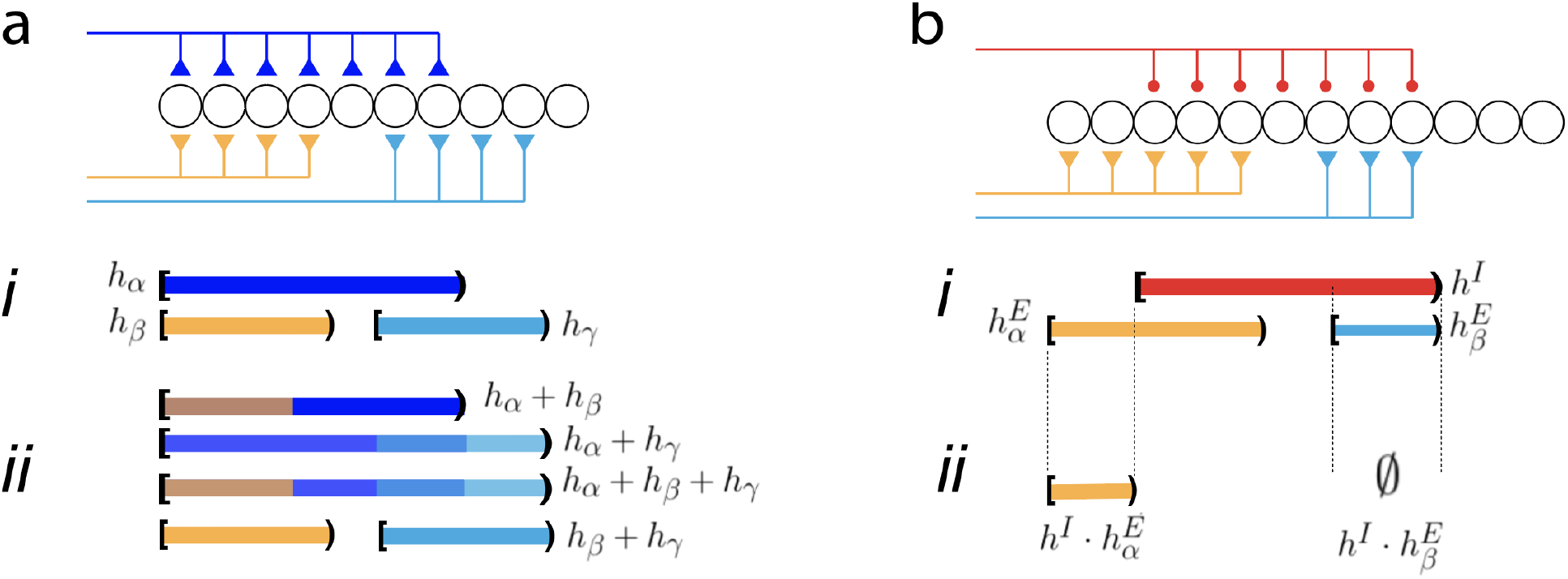
Addition and Multiplication. ***a***, schematic of network receiving three excitatory afferent inputs. ***i***, half-open intervals representing the synaptic fields. ***ii***, addition (union) of different combinations of intervals. **b**, schematic of network receiving two excitatory inputs (orange, cyan) and an inhibitory input (red). ***i***, half-open intervals associated with the activated afferents. ***ii***, multiplication (set minus) of each excitatory intervals by the inhibitory interval.

It is noteworthy that there is some ambiguity from a decoding perspective because the addition operation will fuse overlapping intervals into a larger, single interval (e.g. *h_α_* + *h_γ_* in Fig. 3a). It would not be possible to determine whether a synaptic interval is a result of a single high intensity pure tone, multiple low intensity pure tones with small differences in frequencies, or band limited noise (see Discussion).

Closure is satisfied because the union of an arbitrary number of overlapping intervals results in an interval that is always within the neural space. The union of any number of disjoint intervals is also within the neural space (more accurately, is a subset of the topology defined in neural space, see Supporting Text S1). A notable property that will be important later is that there is no additive inverse that can decrease the interval length.

One example of addition that occurs under biological conditions is when a pure tone arrives simultaneously to the two ears. The signal propagates separately through the auditory pathway but eventually converges at some brain region. Because each input is due to the same tone, the resultant synaptic intervals will be at same location (i.e. have the same starting points) on the tonotopic axis, though their lengths may differ because of interaural intensity differences (orange and blue intervals in Fig. 3a). Fig. 4a (top panel) shows the predicted total length when two intervals with different lengths are added (the length of the cyan interval is fixed while that of the blue is increased). The total length is equal to the length of the longer interval: hence, it is initially constant and equal to that of the cyan interval but then increases linearly when the length of the blue interval becomes longer.

**Figure 4:**
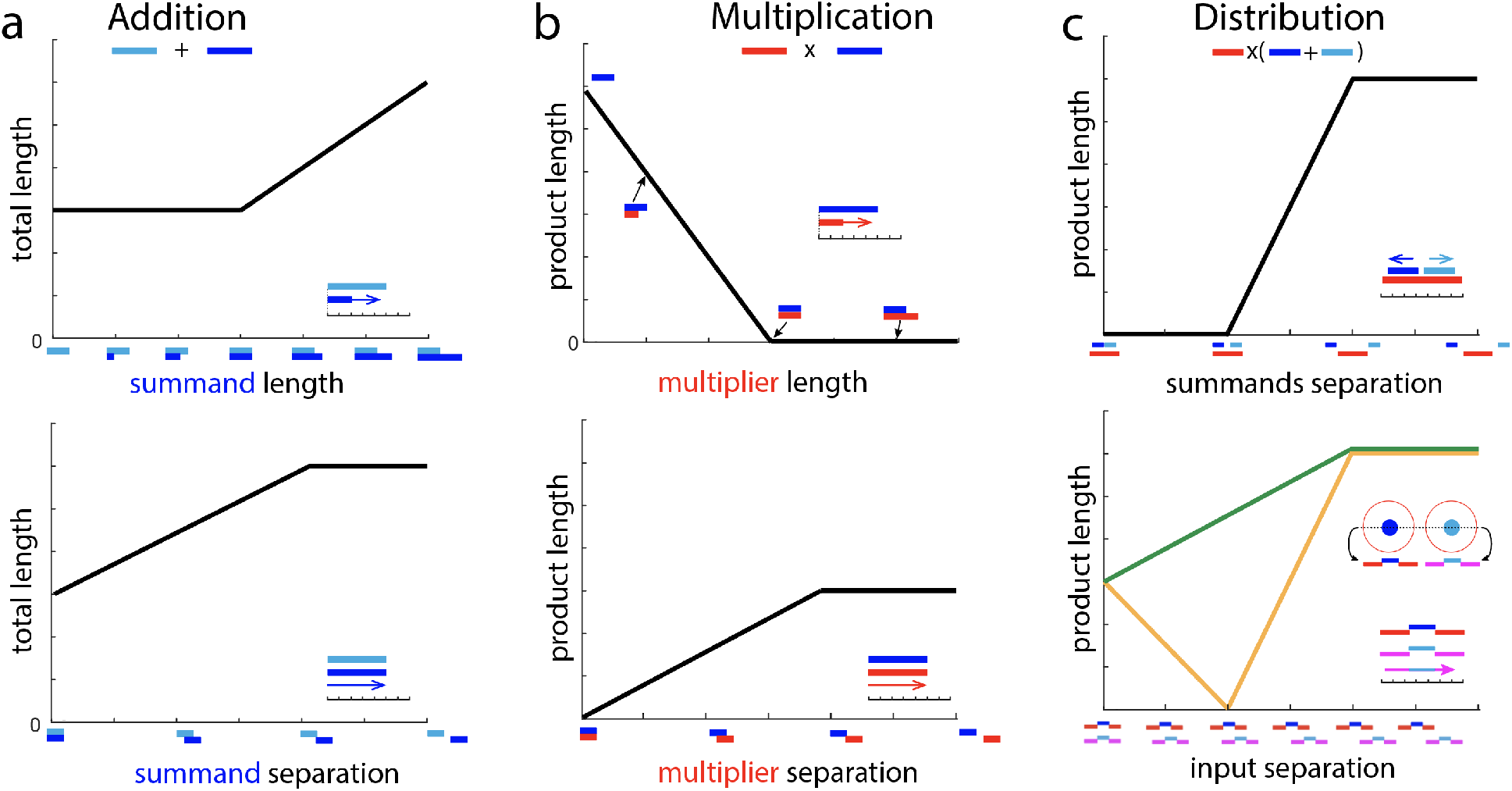
Predicted effects of addition and multiplication. ***a, top***, addition of two intervals (cyan, blue) when the length of one interval (blue) is increased (top). ***bottom***, one interval (blue) is shifted to the right of the other (cyan). ***b*, top**, multiplication of an excitatory interval (blue) by an inhibitory interval of increasing length (red). ***bottom***, multiplication when the inhibitory interval is shifted to the right. ***c*, top**, effects of inhibition (red) on two excitatory intervals (blue, cyan). With the inhibitory interval at a fixed location, the distance between the excitatory intervals is increased systematically. ***bottom***, Effects of two inputs configured as center-surround where the excitatory intervals (blue, cyan) are each flanked by two inhibitory intervals (red, magenta). One of the inputs is shifted systematically to the right of the other. Predicted product length when the inputs occur simultaneously (orange) and when calculated with inputs delivered sequentially.

Addition also takes place when the sound is composed of two pure tones with different frequencies. This would generate two synaptic intervals with different starting points and possibly different lengths (e.g. orange and cyan intervals in Fig. 3a). The total length depends on the degree of overlap between the intervals. In Fig. 4a (bottom panel), the location of one interval (cyan) is fixed while the other (blue) is shifted rightward. When the two intervals completely overlap, the total length is equal to the length of one interval. As the blue interval is shifted, the total length increases linearly and plateaus when the two intervals become disjoint.

Inhibition decreases the excitability of the network and would be expected to reduce the sizeof the synaptic interval. This is not possible with the addition operation because there is no inverse (i.e. ‘subtraction’ is not defined). Therefore, to incorporate the effects of inhibition, a ‘multiplication’ operation is introduced (Fig. 3b). Multiplication (‘·’) of an excitatory synaptic interval *h^E^* by an inhibitory interval *h^I^* is defined as:

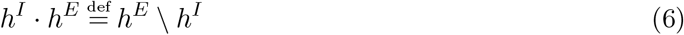

The set minus operation “\” eliminates the points from the multiplicand *h^E^* that it has in common with the multiplier *h^I^* (Fig. 3b*i,ii*) thereby decreasing the multiplicand’s length. Note that this differs from the more common set theoretic definition of multiplication, where the product of sets is given by their intersection. Multiplication yields several results, depending on the relative locations and size of the multiplicand and the multiplier (see Supporting Text S1). If the excitatory (***E***) and inhibitory (***I***) intervals do not overlap, then the ***E*** interval is unaffected. If the intervals overlap (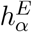 and *h^I^*), then the **E** interval is shortened (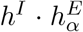 in *ii*). If the ***E*** interval 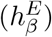 is completely within the ***I*** interval, the product is the empty set, indicating complete cancellation (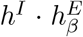 in *ii*). Multiplication can also change the starting points of the intervals and split an interval into two separate intervals (not shown, see Supporting Text S1). Multiplication satisfies the closure property as the resulting interval is a half-open interval that is always in neural space. Other algebraic properties are discussed in Supporting Text S1.

If the excitatory and inhibitory inputs are co-activated (as in feedforward circuits), then the ***E*** and ***I*** intervals will be at the same location (same starting points) on the tonotopic axis but may have different lengths. Fig. 4b (top panel) plots the predicted length of the product when the ***E*** interval is multiplied by ***I*** intervals of increasing lengths. The product length decreases and becomes zero when length of the ***I*** interval exceeds that of the ***E*** interval.

If the ***E*** and ***I*** inputs are independent of each other, the synaptic intervals could be at different locations on the tonotopic axis. Figure 4b (bottom panel) plots the length of the product when the starting point of the ***I*** interval (red) is shifted systematically to the right of the ***E*** interval (blue). The product length is zero when the ***E*** and ***I*** intervals overlap completely (separation = 0) and increases linearly as the overlap decreases, eventually plateauing to a constant value when the intervals become disjoint.

With addition and multiplication defined, the rules for combining the two operations can now be determined. A simple case is when a network receives two excitatory inputs that results in synaptic intervals 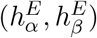 and a single inhibitory input that result in an inhibitory interval (*h^I^*). This scenario would occur if binaural excitatory inputs that converge in a network are then acted on by local inhibitory neurons. When all three inputs are activated simultaneously, the intervals combine in neural space as 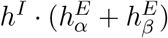. It can be shown that multiplication is *left* distributive so that (see Supporting Text S1):

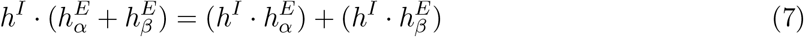

Intuitively, this means that the effect of a single inhibitory input on two separate excitatory inputs can be calculated by computing the inhibitory effects on each separately and then adding the results. Multiplication, however, is not *right* distributive. Thus, given two inhibitory intervals (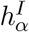 and 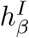) acting on a single excitatory interval (*h^E^*):

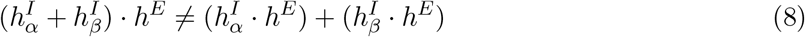

Fig. 4 c (top panel) plots the predicted length when two ***E*** intervals are multiplied by an ***I*** interval as per eq. 8 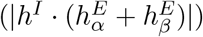. The two ***E*** intervals (blue, cyan) are shifted, respectively, left- and rightward relative to the ***I*** interval (red). The product length is zero as long as the two ***E*** intervals are within the ***I*** interval. When the two ***E*** intervals reach and exceed the borders of the ***I*** interval, the product length increases and reaches a plateau when the ***E*** and ***I*** intervals become disjoint.

A common physiological scenario is when sound is composed of two pure tones and each tone results in an excitatory synaptic field surrounded by an inhibitory field (center surround inhibition, Fig. 4c, bottom panel). The corresponding composite interval contains an excitatory interval that is flanked by two inhibitory intervals (inset). Letting the ***I-E-I*** interval triplet generated by each tone be 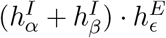 and 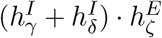, the expression when both occur simultaneously is:

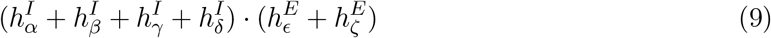

In Fig. 4c (bottom panel, orange curve), the location of one composite interval is shifted rightward. When the composite intervals coincide (separation =0), the product length is equal to that of a single excitatory interval. With increasing separation, the product length decreases towards zero but then increases, reaching a plateau when the excitatory and inhibitory components of each composite interval no longer overlap.

It is worth noting that the effect of introducing two tones simultaneously cannot be predicted by introducing each separately and then combining the results (see Supporting Text S1 for proof). That is,

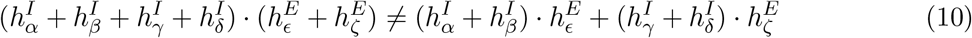

The green curve in Fig. 4 c (bottom panel) is the predicted product length when the ***I -E -I*** triplet pairs are delivered separately and their product lengths subsequently summed. Intuitively, the curves differ because the effects of inhibition on the adjacent excitatory interval is absent; indeed, the result resembles that of adding two excitatory intervals (Fig. 4a, bottom panel). A practical implication is that the intervals due to complex sound cannot be predicted by presenting individual tones separately (see Discussion).

### Simulations with spiking neurons

Key features of the mathematical model were examined with simulations performed on a 2 dimensional network model of spiking excitatory and inhibitory neurons [26]. Both ***E*** and ***I*** neuron population receive a Gaussian distributed excitatory drive from an external source (Fig. 5a); the ***E*** cells in addition receive feedforward inhibitory inputs from the ***I*** cells. Stimulation evokes Gaussian distributed excitatory inward currents in both populations and also inhibitory currents in the ***E*** cells (profiles of currents shown in insets). With brief stimuli, the recurrent connections between neurons [27] do not contribute significantly to activity in auditory cortical circuits [26] and were omitted. The region encompassing neurons that fire is henceforth referred to as the activated area (Fig. 5b, top panel). The underlying synaptic field (bottom panel) is described by the area of the network where the *net* synaptic inputs to cells exceeded (were more negative than) rheobase, the minimum current needed to evoke an action potential (*I_Rh_*, inset). As defined, the synaptic field is a composite of all the inputs, both excitatory and inhibitory, that are evoked during a stimulus. Both the activated area and the synaptic field are quantified either by the diameters of circles fitted to the boundary points (magenta) or by the length of their projections to one axis (orange bar). Note that the spatial dimensions have units of *cell number* (see Methods to convert to microns).

**Figure 5:**
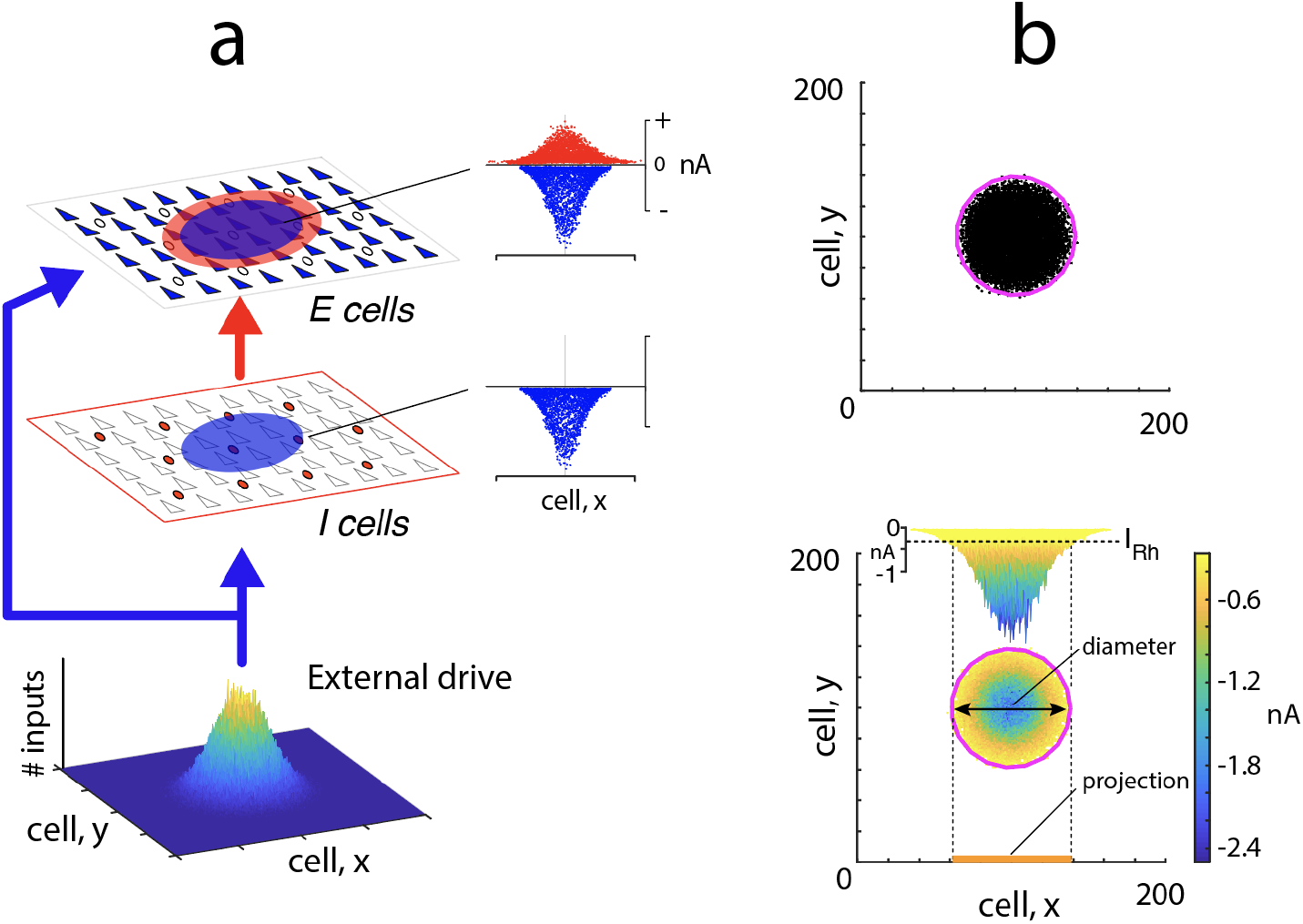
Simulations with spiking neurons. ***a***, Network consists of excitatory and inhibitory neuron populations. An external drive evokes excitatory inputs in both populations (blue disks) and inhibitory inputs to the ***E*** cells (red disk). ***insets***, profiles of excitatory (blue) and inhibitory (red) currents evoked in the ***E*** and ***I*** populations. ***b, bottom***, synaptic field evoked in the network during a stimulus. The spatial extent of the synaptic field is quantified either by the diameter of a circle fitted to its outermost points (magenta) or by the length of its projection to the tonotopic axis (orange bar). ***inset***, profile of net synaptic current generated in the ***E*** cell population. The perimeter of the synaptic field encompasses cells whose net synaptic current input exceeded rheobase (*I_Rh_*). ***top***, activated area contains cells that fired action potentials (dots).

To test the addition operation, two external excitatory drives were delivered to the center of the network simultaneously (without inhibition). Increasing the width (by increasing the standard deviation *σ_α_* of the external drive) of one stimulus, while keeping that (*σ_β_*) of the other fixed, increased the diameters of the synaptic field and activated areas (Fig. 6a, top panel, *i-iii*). As predicted, the diameters of the synaptic field (bottom panel, orange) and activated area (black) initially did not change but then increased as *σ_α_* continued to widen. However, because the synaptic currents were Gaussian distributed (Fig. 5b, bottom panel), the curve started to increase before *σ_α_* became equal to 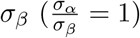. When delivered simultaneously, the magnitude of the composite current increased, causing the region that exceeded rheobase to widen (Fig 6 a, top panel, compare synaptic field evoked with a single stimulus (*i*) to that evoked with 2 stimuli (*ii*)). The diameter can be calculated from the standard deviations of the two inputs (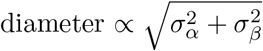, not shown).

**Figure 6:**
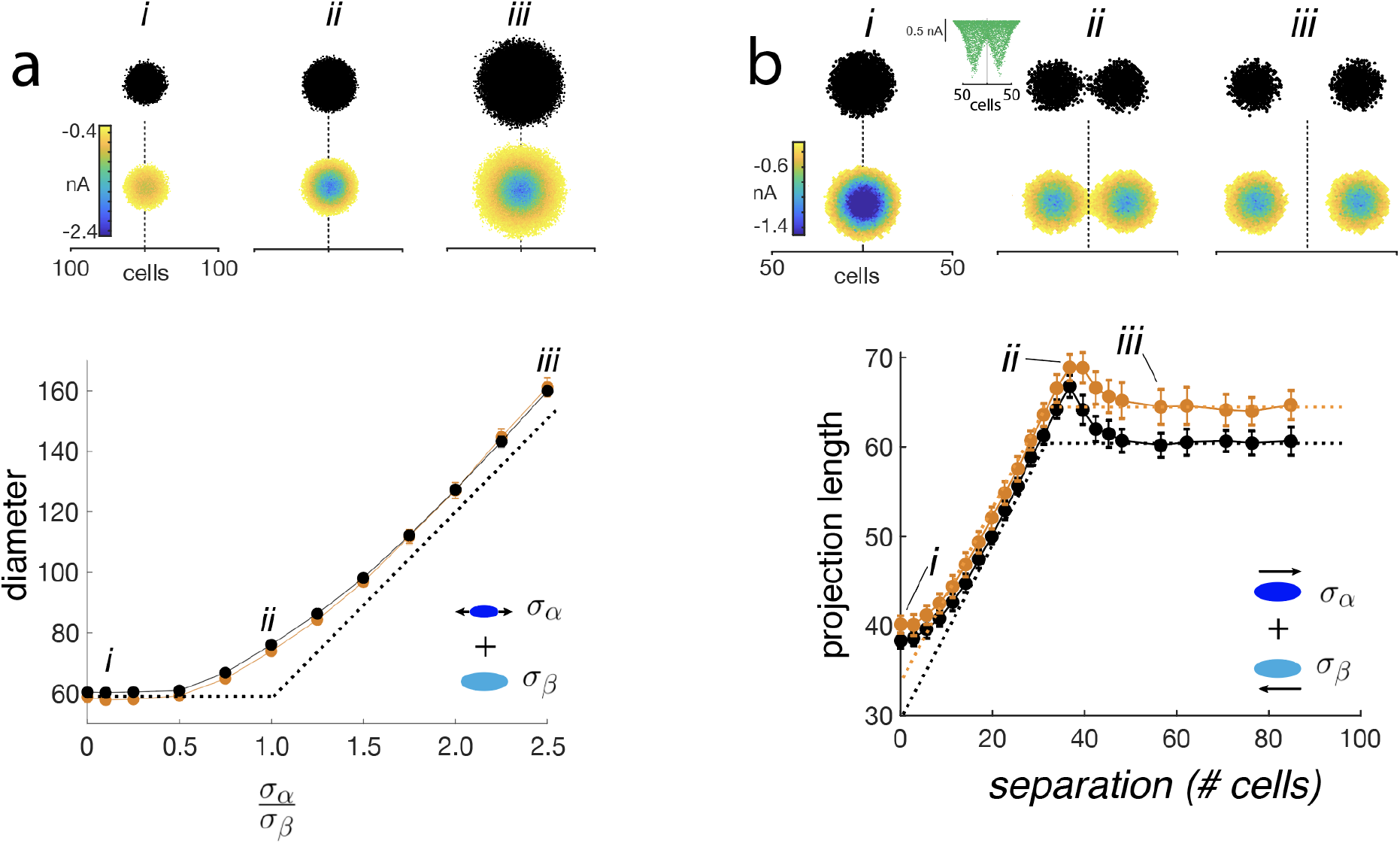
Test of addition. ***a, top***, Activated area (top; 1 sweep) and underlying synaptic field (bottom; average of 25 sweeps). ***i***, one stimulus. ***ii-iii***, two stimuli delivered to the center of the network. The width of one input was systematically increased (*σ_α_*: 2-45 cells) while that of the other (*σ_β_* = 20 cells) was kept constant. ***bottom***, plot of synaptic field (orange) and activated area (black) diameters vs 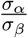. Dashed curve is predicted relation. ***b***, addition of two spatially separated excitatory inputs (*σ_α_* = *σ_β_* = 10). ***top, i - iii***, activated areas and synaptic fields with increasing stimulus separation. Inset in ***ii*** shows example of excitatory synaptic current profiles. ***bottom***, projection length vs. separation distance for synaptic field (orange) and activated area (black). Dashed curves are predicted changes.

To examine the addition of spatially disparate synaptic fields, two excitatory inputs were delivered at different distances from each other (Fig. 6b, top panel). Consistent with the prediction, the projection lengths of the synaptic field (bottom panel, orange) and activated areas (black) increased with stimulus separation and reached a plateau when the two inputs became disjoint (*iii*). The projection lengths were greater than predicted (dashed lines) when the separation was small (< 10 cells) and when the intervals were just becoming disjoint (at separation ~ 40 cells, *ii*) due to the summation properties of the Gaussian distributed inputs discussed above.

To test the multiplication operation, the ***E*** and ***I*** neurons were stimulated simultaneously, resulting in excitatory and inhibitory synaptic currents in the ***E*** cells (inset in top panel of Fig. 7a *ii*). The width of the excitatory input (*σ*_exc_) was kept constant while that of the inhibitory input (*σ*_inh_) was increased systematically. As predicted, the diameter of the synaptic field (bottom panel, orange) and activated area (black) decreased with increasing *σ_inh_*. However, the diameter asymptoted towards a non-zero value. Because the network was feedforward, the inhibitory input was delayed relative to excitation by about 10-15 ms; as a result, there was always a time window where excitation dominated. The excitatory synaptic input was not canceled even when the inhibition was twice as wide (top panel, *iii*).

**Figure 7:**
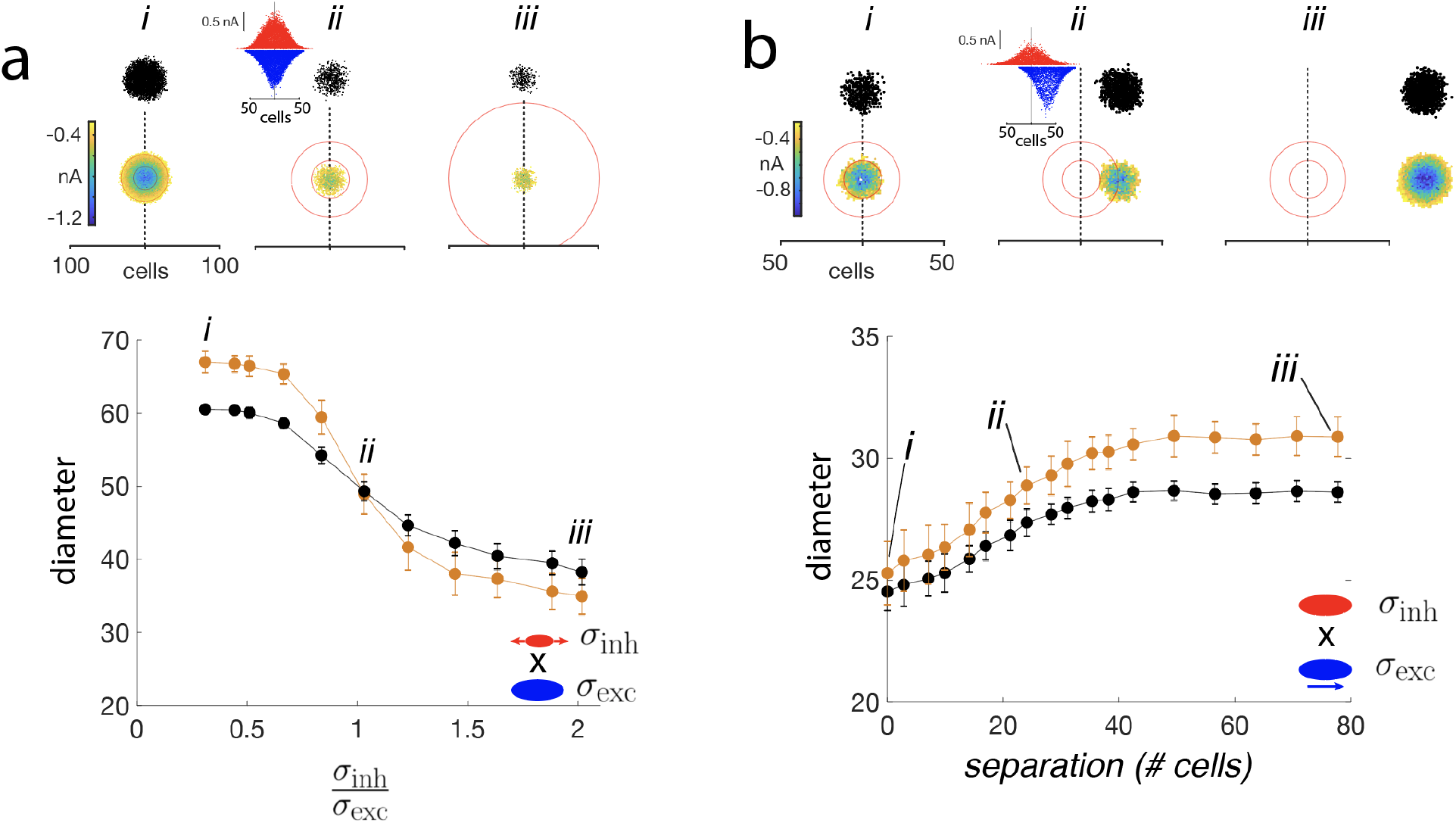
Test of multiplication. ***a, top, i-iii*** activated areas and synaptic fields evoked with excitatory (*σ_exc_* = 20) and inhibitory (*σ_inh_* = 2 – 45) inputs. The spatial extent of the inhibition is demarcated by the red circles (inner circle: 1 *σ_inh_*; outer: 2 *σ_inh_*). Inset in ***ii*** shows an example of excitatory (blue) and inhibitory (red) synaptic current profiles. ***bottom***, plot of activated area (black) and synaptic field (orange) vs 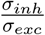. ***b***, Same as in ***a*** except that the excitatory input (*σ_exc_* = 10) was shifted systematically to the right of inhibition (*σ_inh_* = 10). ***bottom***, Diameters of activated area (black) and synaptic field (orange) plotted against separation between excitatory and inhibitory synaptic fields

To examine multiplication of spatially disparate ***E*** and ***I*** inputs, the excitatory input was shifted systematically to the right of the inhibitory input (Fig. 7b). As predicted, the diameters of the synaptic field (bottom panel, orange) and activated area (black) increased with the ***E -I*** separation and plateaued when the ***E*** and ***I*** inputs became disjoint (*iii*).

To examine how multiplication distributes over addition, two excitatory inputs and one inhibitory input were delivered simultaneously to the network (Fig. 8a). This is the analog of the left hand side of (eq. 8). All three inputs were initially at the center and then with the inhibition stationary, the two excitatory inputs were shifted left and right (Fig. 8a, top panel, *i-iii*). As predicted, the projection length of the synaptic field increased towards an asymptotic value (bottom panel, orange). To reproduce the right hand side of eq. 8, simulations were performed with inhibition, first with one of the excitatory inputs and then with the other; the resultant projection lengths of each were then summed (green). As was observed with simultaneous stimulation, the projection length increased with separation. The match was poor at small separations <10 cells (*i*) and at separation of ~ 30 cells (*ii*) because the interaction between the Gaussian excitatory currents (see above) did not factor in when each input was delivered separately. The two curves were nearly identical at separations of 40-80 cells. In this range the ***E*** inputs were disjoint (as indicated by the plateauing of the excitation-only curve (dashed orange)) but still overlapped with the inhibitory synaptic field (the orange and green curves were below the excitation-only curve).

**Figure 8:**
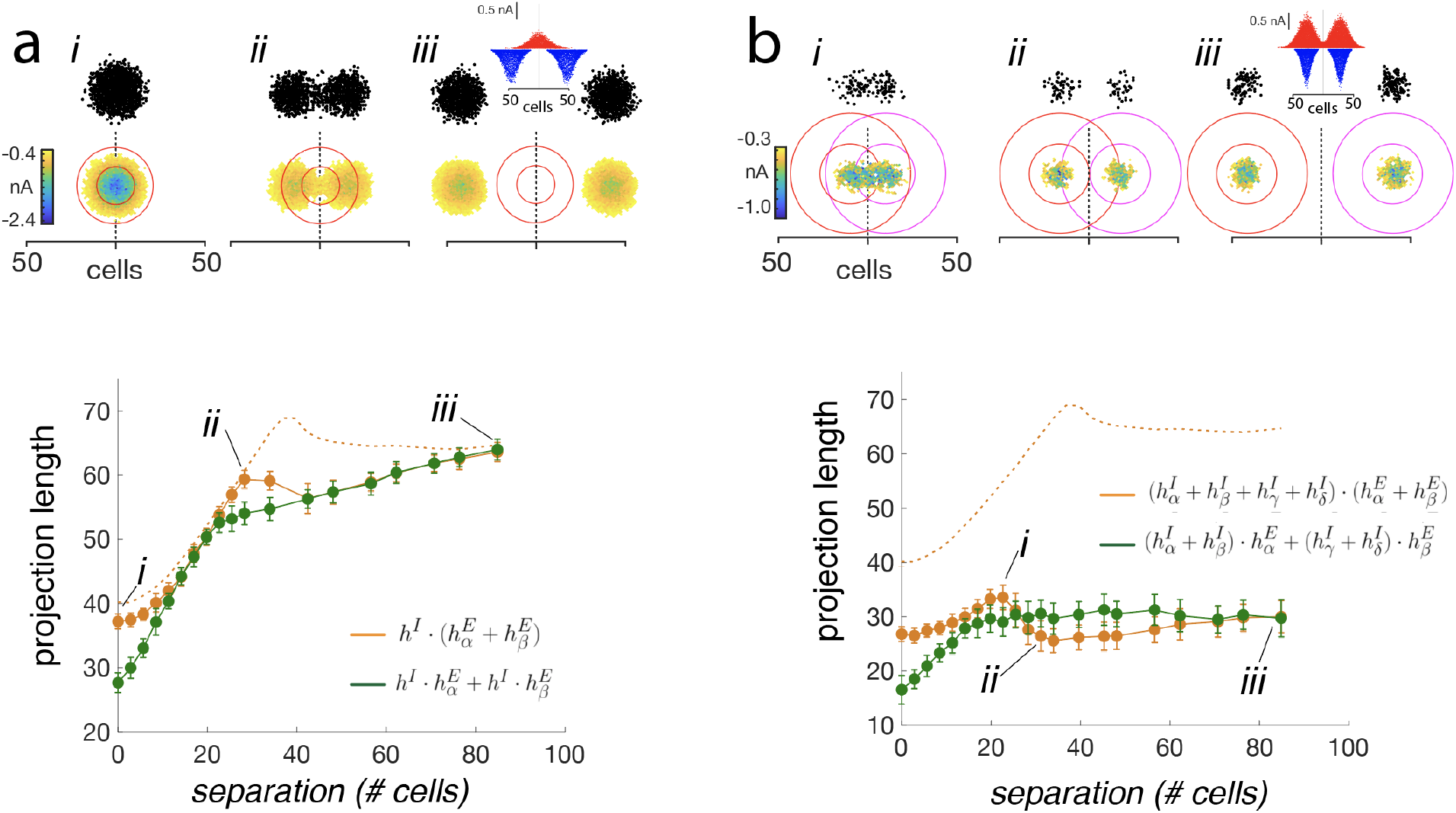
Test of distributive properties. ***a, top, i-iii*** Representative activated areas and synaptic fields generated by two excitatory inputs and a single inhibitory input (*σ_exc_* = *σ_inh_* = 10). With the location of inhibition fixed, the two excitatory inputs were separated systematically. The spatial extent of the inhibition is demarcated by the red circles (inner circle: 1 *σ_inh_*; outer: 2 *σ_inh_*). Inset in ***iii*** shows the excitatory (blue) and inhibitory (red) synaptic current profiles. ***bottom***, plot of synaptic field projection lengths (orange) vs separation of the excitatory inputs. Green symbols are projection lengths obtained with the sequential stimulation protocol (see text). Dotted curve is with no inhibition. ***b***, Simulations with two excitatory-inhibitory pairs, each with center-surround configuration (see inset in ***iii***). ***bottom***, legend as in ***a*** except that the green traces plot the projected lengths obtained when each input (*σ_exc_* = 10, *σ_inh_* = 17) was delivered sequentially (see text).

Finally, the interaction of inputs with center-surround inhibition (eq. 9) was examined by delivering two excitatory inputs, each with associated inhibitory components (inset in Fig, 8b, top panel, *iii*), to the network. The distance between the inputs were then increased systematically and the projection lengths measured (bottom panel, orange curve). At separations > 20 cells, the projection length of the synaptic field decreased to a minimum (*ii*) and then increased towards a plateau (*iii*), consistent with the prediction (Fig. 4c, bottom panel). However, at separations < 20 cells, the length increased instead of decreasing; this is likely due to the interaction of the Gaussian distributed excitatory fields discussed above. To confirm that the same result cannot be obtained by presenting each stimulus separately (right hand side of eq. 10), each ***E -I*** pair was delivered sequentially and the individual projection lengths summed. Unlike with simultaneous stimulation and consistent with the prediction (green curve in Fig. 4c, bottom panel), the projection length increased monotonically to a plateau without a dip (green curve).

## Discussion

The aim of this study was to develop a mathematical framework for a place-code and derive the underlying principles for how tones of varying frequencies and intensities are represented, assembled, and modulated in networks of excitatory and inhibitory neurons. The analyses are not intended to replicate the detailed aspects of biological networks but rather to clarify the minimal conditions that must be met for a viable place coding scheme, to aid in the interpretation of experimental data, and to provide a blueprint for developing computational models. The formal approach ensures that the terms of the model are defined precisely and the proposed computations are constrained strictly by well-established mathematical rules.

### Place code framework in auditory processing: evidence and implications

The model has several implications with regards to auditory processing. In this section, the advantages of the place coding framework are discussed and experimental data are interpreted within the context of the mathematical framework. The formal approach is then applied a well-known psychophysical phenomena to elucidate the underlying computations and network mechanisms.

### Representation of frequency and sound pressure

A key feature of the model is that the ‘functional unit’ of neural space is a set of contiguous neurons that have flexible borders. The associated mathematical architecture, or topology, is a collection of half-open intervals of varying lengths. The model provides a framework for encoding both frequency and intensity (or sound pressure) with a purely place-coding scheme. This is advantageous for brief stimuli where firing rate and spike timing [7, 25, 11] may not be available (see Introduction). Frequency and intensity discrimination does improve with stimulus duration, suggesting that the other variables play complementary roles in improving coding and perception [8, 9, 10, 11, 28].

A network with flexible functional units is also advantageous for maintaining both high resolution frequency and pressure representations. This can be appreciated by comparing the resolutions attainable with the classical columnar organization [18, 20] (the stimulus is assumed brief so that firing rate information is unavailable; see Introduction). In this scheme, the neural space is divided into non-overlapping columns with fixed dimensions and distinct borders. The frequency of a stimulus is encoded by the location of the active column and sound pressure by the number of active neurons within the column (i.e. population rate code). The disadvantage is that there is a trade-off between the achievable frequency and sound pressure resolutions. Intuitively, to maximize the number of frequency levels, the columns should be as small as possible so that more can fit along the tonotopic axis; however, this reduces the number of pressure levels that can be encoded because there are fewer neurons within a column. As shown in Supporting Text S1, the number of frequency levels that can be represented is inversely proportional to the number of pressure levels 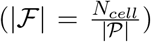, where *N_cell_* is the number of cells in the network. In contrast, for a network with flexible borders, the relation is linear: 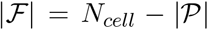. Thus, compared with columns, fewer neurons are needed to have an equal number of frequency and pressure levels.

The advantage of a columnar organization is that the components of a multi-frequency stimulus remain separated in neural space. With flexible units, two intervals generated by two tones with small frequency differences and/or high intensities can fuse into a single interval. In such a case, the inverse mapping (*ψ*^−1^) yields a single tone. As discussed below, ambiguities in perception of complex stimulus are more consistent with a flexible unit organization.

### Relation to difference limens

The Δ*f*, which is used to discretize acoustic space into equivalence classes, is loosely related to minimum discriminable frequency difference measured experimentally (frequency difference limen or *fDL*) [8, 9, 10, 11]. As defined, Δ*f* is a constant but the measured *fDL* will be 0 < *fDL* ≤ Δ*f*, depending on where the frequencies (*f_α_, f_β_*) of the test tones fall within an equivalence class. For example, given two adjacent equivalence classes [*f*_0_, *f*_0_ + Δ*f*) and [*f*_1_, *f*_1_ + Δ*f*), *fDL* will be small if *f_α_* and *f_β_* are close *f*_0_ + Δ*f*, near the border: initially, both *f_α_* and *f_β_* are classified (or ‘heard’) as *f*_0_ but just a small increase in *f_β_* will put it in the next equivalence class and be classified as *f*_1_. If the two frequencies are closer to *f*_0_, a larger increase in *f_β_* is needed for the jump to occur. In principle, this prediction can be tested by presenting two indiscriminable test tones with small differences in frequencies (*ϵ* << Δ*f*); increasing both frequencies with e constant should cause the tones to be discriminable at intervals of ~ Δ*f*.

### Addition operation

The addition operation applied to synaptic intervals is defined as their union: 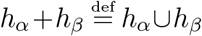. An important consequence is that if the intervals overlap, they will fuse into a single, longer interval. Under physiological conditions, this would occur if tones of a multifrequency stimulus have small differences in frequencies. This is in line with psychophysical experiments, which show that subjects perceive tones with small differences in frequencies as a single tone [29, 30] and have difficulties distinguishing the individual components of a multi-frequency stimulus [31, 32].

Another consequence is that addition of two overlapping intervals is sublinear: |*h_α_*| + |*h_β_*| > |*h_α_* + *h_β_*|. If one interval is also a subset of the other (*h_α_* ⊂ *h_β_*), then the sum is equal to the larger of the two intervals: |*h_α_*| + |*h_β_*| = |*h_β_*|. This scenario would occur when binaural inputs converge onto a common site. Consistent with the prediction, electrophysiological recordings from neurons in inferior colliculus show that the FRAs (assumed to be representative of activity spread, see above) evoked binaurally is equal to larger of two responses evoked monaurally [33] (see Supporting Fig. S1). Similarly, assuming that loudness perception is linked to the length of the interval, a possible psychophysical analog is that a tone presented binaurally to a subject is perceived to be less than twice as loud as monaural stimulation [34]. The apparent sublinear effects may be due to the properties of addition operation, though inhibitory processes are also likely to contribute.

### Multiplication operation and distributive properties

Multiplication of two synaptic intervals is defined as the set minus operation: *h_α_ · h_β_* = *h_β_* \ *h_α_*. The operation removes from the multiplicand (*h_β_*) elements that it has in common with the multiplier (*h_α_*), thereby shortening it. The effect of inhibition can be inferred from the FRAs of neurons. Applying GABA blockers causes the FRAs to widen [35, 36]. If, as mentioned above, the FRA can be used as a proxy for the spatial extent of activated neurons, then the result is consistent with inhibition shortening the synaptic intervals.

The manner in which multiplication distributes over addition has important implications for combining information from multiple sources. In auditory cortex, excitatory pyramidal neurons receive convergent afferent inputs from the thalamus and other pyramidal cells [37, 38]. The two afferents also appear to innervate a common set of local inhibitory neurons [27, 38]. The fact that multiplication is left distributive (eq. 7) means that the effect can be estimated by measuring the effects of inhibition (*h^I^*) on each excitatory inputs 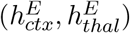 separately and then summing the results: 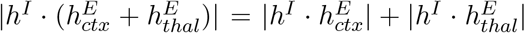. However, because multiplication is not right distributive (eq. 8) a similar approach cannot be used to examine two sources of inhibition acting on a single excitatory interval. In cortical circuits, for example, excitatory neurons can be innervated by several classes of local inhibitory interneurons [39].

More generally, the representation of complex sound with a place coding scheme cannot be predicted by combining the representations of individual components if the inhibition generated by each component interact. As shown in Eq. 10 and Fig. 4c (bottom), the response of two tones presented simultaneously is not a simple combination of the responses to each tone separately. This is in line with experimental data showing that the response of neurons to pure tones and white noise stimuli differ [40]. It should be emphasized that this conclusion was derived mathematically from the distributive properties; it not trivially related to non-linearities contributed by e.g. inhibitory conductances or voltage gated channels since the model has no biophysical variables.

### Transmission of acoustic information

At a more abstract level, the mathematical structures of the place coding scheme set the rules for the transfer of signals across brain regions along the auditory pathway. For example, faithful transmission of frequency and pressure information might require that the mapping be homeomorphic, which would necessitate that the topology of the source and target brain regions be the same. If the algebraic operations also need to be preserved, then the mapping should also be a homomorphism. The mathematical conditions set biological constraints on the circuitry and connection patterns of the brain regions and can be used a guide for experiments.

#### Algebra of loudness summation

In the auditory system, the perception of simultaneously presented tones or band limited noise depends on whether the frequencies are within the so-called critical band (CB) of frequencies [41, 42]. Figure 9a shows an experiment where subjects were asked to judge the loudness of a band limited white noise [43, 44]. As the bandwidth is increased, the perceived loudness is initially constant but then increases when the bandwidth exceeds CB (vertical line). Increasing the intensity of the stimulus shifts the curves up but does not change CB. The origin of the CB is unclear and there is debate as to whether it is peripheral involving mainly excitatory processes [45, 46], or central, which may also recruit inhibition [47, 48, 49]. The tonotopic axis is often divided into 24 CBs, each uniquely identified by the center frequency [42]. In the following, algebraic operations are used to describe features of loudness summation and to suggest network mechanisms.

**Figure 9:**
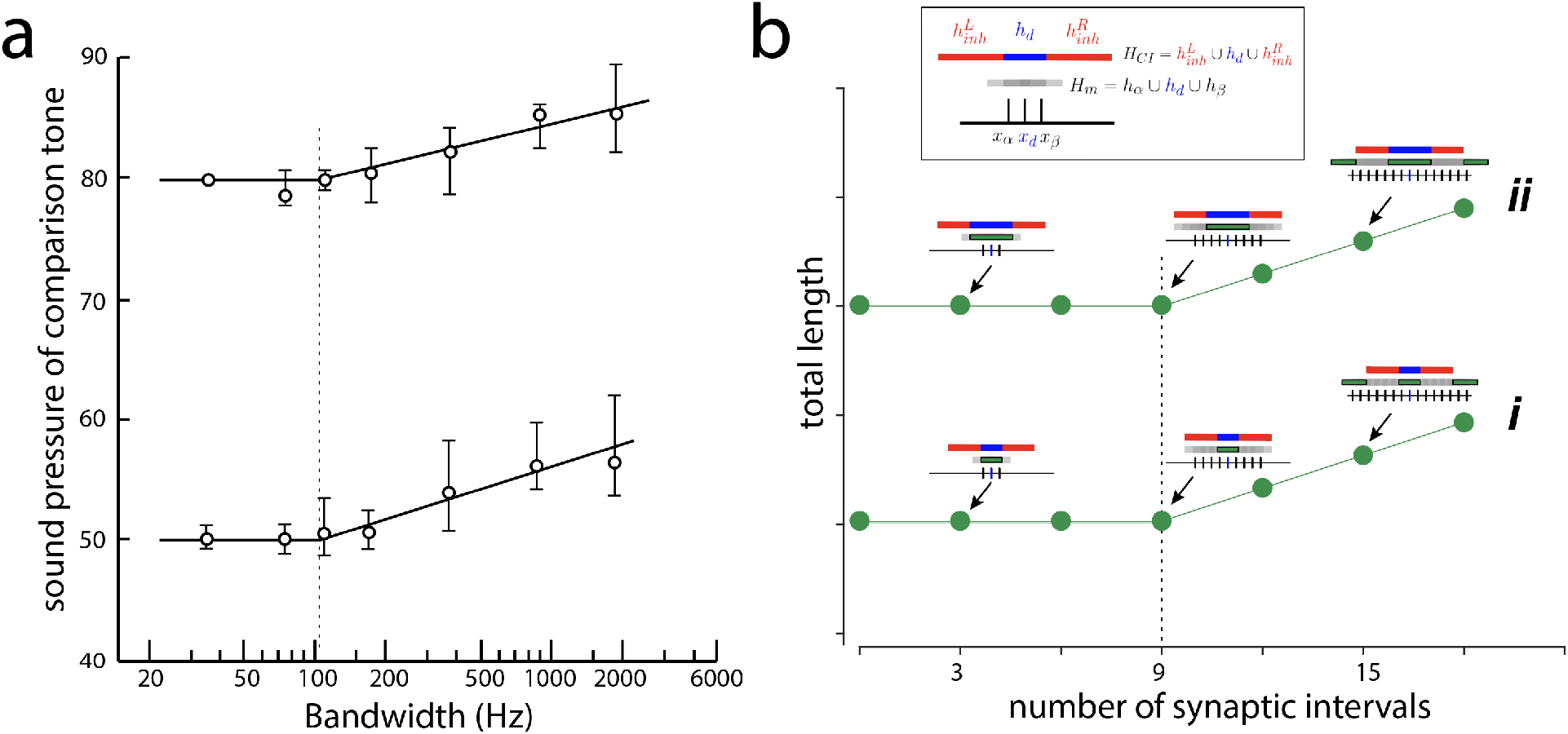
Algebra of loudness summation. ***a***, Perceived loudness when subjects were presented with white noise of increasing band width (abscissa) at 2 sound pressure levels (lower and upper traces). Subjects were asked to increase the intensity of a pure tone until it was as loud as the band-limited white noise (ordinate). Dotted vertical line corresponds to critical band. Figure adapted from [44]. ***b***, Predicted interval lengths resulting from the interaction of multi-tone stimulus delivered simultaneously. ***Boxed inset***, overlapping synaptic intervals (*H_m_*, gray) generated by stimulus with 3 frequency components. Tic marks show location of interval centers (*x_α_, x_d_, x_β_*) along tonotopic axis. The dominant tone (blue) also generates two inhibitory side bands (red). Plot shows resultant length (*h_l_*) after the operations (see text) as the number of intervals in *H_m_* is increased (abscissa). Green bars in insets show portion of *H_m_* that was not cancelled by inhibition. Dotted vertical line marks deviation from constant value.

A band-limited noise stimulus, or more generally a complex stimulus with multiple tones, may be expressed, after discretization, as a set of sequential frequencies: *F_m_* = {*f_i_* | *i* = (1, 2,…, *n*); *f*_*i*+1_ – *f_i_* = Δ*f*). The ‘bandwidth’ is defined as the difference between the highest and lowest frequency components (*BW* = *f_n_* – *f*_1_). In neural space, the stimulus results in a set of excitatory intervals that is the union of individual excitatory intervals generated by each tonal component: 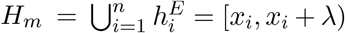, where λ is the length of each interval and is the same for all intervals. Note that because the frequencies for a band limited noise stimulus is a contiguous set, *H_m_* is a single interval: *H_m_* = [*x_i_, x_n_* + λ).

The model assumes that in the multi-tone stimulus, one of the tones with frequency *f_d_* ∈ *F_m_* is dominant and generates inhibitory intervals (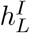 and 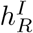) that abut the excitatory interval 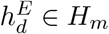 with no overlap 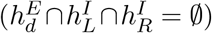, as in a so-called lateral inhibitory configuration (see Suppl. text for formal definitions). The other tones may also generate inhibitory intervals but they should be much smaller. Physiologically, the dominant tone may be the one with the lowest frequency, which has been shown to mask higher frequency components [30], or defined as the tone at the center of a CB [42]. The boxed inset in Fig. 9b shows the relationship between *H_m_* (gray, composed of 3 intervals), 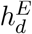 (blue), and the two inhibitory intervals (red). The length of the interval *h_l_* that results from the interaction of these 4 intervals is taken to be a proxy for loudness perception and is described by (see Suppl. text for derivation and proofs):

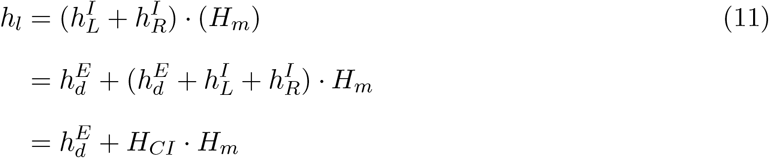

where 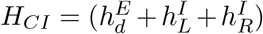 is henceforth termed the critical interval. Intuitively, the second term on the right hand side means that portions of *H_m_* that overlap with critical interval are cancelled by multiplication. The first term on the right is the synaptic interval generated by the dominant tone, which is part of *H_m_* and does not overlap with the inhibition. Fig. 9b shows the result graphically when *H_m_* is lengthened by adding more tones to the stimulus. The length is constant 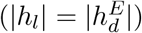 until the number of components is such that *H_m_* exceeds the boundaries of the critical interval (dotted vertical line). In this example, the deviation occurs when the number of intervals, and hence the number of frequency components, exceed 9. The CB is then *f*_9_ – *f*_1_.

Increasing the intensity of the stimulus amounts to increasing the lengths of the excitatory synaptic intervals. As shown in the supplementary text, the CB will not change provided the inhibitory interval lengths do not change. Under these conditions (see Supporting text), lengthening of intervals only affects the first term of eq. 11, resulting in an increase in baseline (upward shift of curves) without a change in CB (compare curves ***i,ii*** in Fig. 9b).

An all-excitatory version without inhibition will not reproduce the data: the critical interval would then be 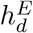 and since 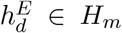, *h_l_* will be greater than 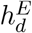 if *H_m_* has more than one component and will grow with increasing number of tonal components. Unlike the data, the curves would have no flat region. An intermediate version is to weaken the inhibition so that the intervals in *H_m_* are merely shortened rather than completely cancelled. The critical interval is still 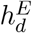 but because the other intervals are reduced in length (less than 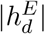), *H_m_* can have more than one interval and still be within the critical interval. However, the CB will be narrower than with strong inhibition.

The operations have also been used to describe a related experiment where instead of white noise, 4 tones with equally spaced frequencies were used as stimuli. Increasing the frequency difference (spacing) between the tone produced curves similar to those of Fig. 9a [44] (see supplementary Fig. S2).

Equation 11 is not intended to suggest specific circuit mechanisms but rather to elucidate the general requirements for loudness summation. While there is some evidence for a dominant tone [30] and inhibitory processes [49], the extent of e.g. inhibitory intervals is less clear and is likely to reflect the combined effect of the individual excitatory and inhibitory intervals generated by other tones in the stimulus. The precise mechanisms needs to be systematically explored with more detailed analyses, simulations, and experiments. The two assumptions-the existence of dominant tone and an associated inhibitory interval that doesn’t vary with intensity-need to be verified experimentally.

### Assumptions and limitations

The mathematical model is based on two salient features of the auditory system. One is that the neural space is organized tonotopically. Tonotopy has been described in most neural structures in the auditory pathway, from brainstem areas [4, 50, 51] to at least layer 4 of primary auditory cortex. Whether tonotopy is maintained throughout cortical layers is controversial, with some studies (all in mice) showing clear tonotopy [52, 53, 54, 55] and others showing a more ‘salt-and-pepper’ organization [56, 57, 55]. A salt-and-pepper organization suggests that the incoming afferents are distributed widely in the neural space rather than confined to a small area. The model needs a relatively prominent tonotopy to satisfy the requirement that synaptic intervals encompass a contiguous set of cells.

A second requirement is that the size of the synaptic interval and activated area increase with the intensity of the sound. Intensity-related expansion of response areas occurs in the cochlea [23, 24, 58] and can also be inferred from the excitatory frequency-response areas (FRAs) of individual neurons. The excitatory FRAs, which document the firing of cells to tones of varying frequencies and intensities, are typically “V-shaped” (see Supporting Figure S1 for examples). At low intensities, neurons fire only when the tone frequencies are near its preferred frequency (tip of the V). At higher intensities, the range of frequencies that evoke firing increases substantially [59, 53]. If adjacent neurons have comparably-shaped FRAs but have slightly different preferred frequencies, an increase in intensity would translate to an increase in the spatial extent of activated neurons.

For mathematical convenience, the location of the synaptic intervals was identified by the leftmost point (closed end) of the interval, with increases in intensity signaled by a lengthening of the interval in the rightward (high frequency) direction. Similar behavior has been observed in the cochlea albeit in the opposite direction: an increase in the intensity causes response area to increase towards low frequency region of the basilar membrane while the high frequency cutoff remains fixed [58, 24, 3]. The choice of the leftmost point to tag the interval is arbitrary and any point in the interval will suffice provided an analogous point can be identified uniquely in each interval in the set. Experimentally, using the center of mass of active neurons as the identifier might be more practical.

For simplicity, both Δ*f* and Δ*p* are kept constant along the tonotopic axis, which is inaccurate because the range of frequencies and sound pressure changes with frequency and sound pressure level. To represent the full ranges, the frequency and pressure can be transformed into an octave and decibel scales prior to mapping to neural space.

The algebraic operations were derived from set theoretic operations and the magnitude of the underlying synaptic inputs were irrelevant. Under biological conditions, the input magnitude determines the degree to which biophysical, synaptic, and network processes become engaged, which will affect the length of the synaptic intervals and activated areas. Not surprisingly, the results of the network simulations deviated quantitatively from the mathematical predictions in some regimes (compare Fig. 4 to Figs 6, 7). Most of the discrepancies in the simulations were because the magnitudes of the synaptic inputs were Gaussian distributed along the tonotopic axis. In biological networks, the discrepancies may be exacerbated by the presence of threshold processes such as regenerative events [60, 61]. The underlying algebraic operations may be obscured in regimes such as these.

The model incorrectly assumes that the strength of inhibition is sufficiently strong to fully cancel excitation. This facilitated analysis because the effect of multiplication depends solely on the overlap between the multiplicand and multiplier. As the simulations with the feedforward network showed, the excitation cannot be fully canceled by inhibition owing to synaptic delay. Moreover, the balance may be spatially non-homogeneous: in center-surround suppression, excitation dominates at the preferred frequency with inhibition becoming more prominent at non-preferred frequencies [59, 35, 36]. To apply multiplication to biological systems, it may be necessary to define empirically an “effective” inhibitory field that takes into account for ***E -I*** imbalances.

Though the simulations that were used to test the analyses predictions used a network model based on cortical circuits [26, 27], the results should generalize to other network types provided the stimuli are brief (50 ms) so that cells fire only a single action potential. The mathematical model treats neurons as binary units and so only the first action potential is important. Hence, if the stimulus is brief and suprathreshold, the results obtained with a network consisting of e.g. repetitively firing cortical neurons [14, 27] or transiently firing bushy cells [62] will be qualitatively similar. The results are likely to differ with longer duration stimuli, which would allow various time- and voltage-dependent channels to become active and engage recurrent connections.

## Methods and Materials

Simulations were performed with a modified version of a network model used previously [26]. Briefly, the model is a 200 x 200 cell network composed of 75% excitatory (***E***) and 25% inhibitory (***I***) neurons. The connection architecture, synaptic amplitudes/dynamics, and intrinsic properties of neurons were based on experimental data obtained from paired whole-cell recordings of excitatory pyramidal neurons and inhibitory fast-spiking and low threshold spiking interneurons [27]. For this study, the low-threshold spiking interneurons and the recurrent connections between the different cell types were removed, leaving only the inhibitory connections from fast spiking interneurons to pyramidal neurons. The connection probability between the inhibitory fast-spiking cells and the excitatory pyramidal cells was Gaussian distributed with a standard deviation of 75 μm and peak of 0.4 [27].

Both ***E*** and ***I*** cells received excitatory synaptic barrages from an external source. The synaptic barrages to each cell (50 ms duration) represented the activity of a specified number of presynaptic neurons. The average number (*n_in_*(*x, y*)) of inputs that each neuron at location *x,y* received followed a Gaussian curve so that cells at the center of the network received more inputs (Fig. 5 a, bottom). For each run, the number was randomized by drawing a number from a Gaussian distribution with mean *n_in_*(*x, y*) and a standard deviation 0.25 * *n_in_*(*x, y*) so that the synaptic fields and activated areas varied from trial to trial. Excitatory synaptic currents were evoked in the ***E*** and ***I*** cell populations and inhibitory synaptic currents in the ***E*** cell population after the ***I*** cells fired (insets in Fig. 5a). The spatial extents of the synaptic inputs were varied by changing the standard deviations of the external drive. In some simulations, the ***E*** and ***I*** cell populations were uncoupled and received separate inputs that could be varied independently of each other.

The neurons are adaptive exponential integrate-and-fire units with parameters adjusted to replicate pyramidal and fast spiking inhibitory neuron firing (see [26] for the parameter values). The cells fired at most one action potential when stimulated with the synaptic barrage.

The synaptic field was defined as the area of the network where the net synaptic currents to the cells exceeded rheobase, the minimum current needed to evoke an action potential in the ***E*** cells (*I_Rh_*, inset in Fig. 5b, bottom panel). *I_Rh_* was estimated by calculating the net synaptic current near firing threshold (*V_θ_*): *I*_net_ = *g_exc_* * (*V_θ_* – *E_exc_*) + *g_inh_* * (*V_θ_* – *E_inh_*) where *g_exc_, g_inh_* are the excitatory and inhibitory conductances, respectively, and *E_exc_* = 0 mV, *E_inh_* = −80 mV are the reversal potentials. For the ***E*** cells, rheobase is approximately −0.27 nA.

The spatial extent of the synaptic field or activated area was quantified as the diameter of a circle fitted to the outermost points (maroon circles in Fig. 5b). In simulations with multiple components, the spatial extents were quantified as the total length of the projection onto the tonotopic axis (orange bar in Fig. 5b, bottom panel). The diameters and lengths have units of cell number but can be converted to microns by multiplying by 7.5 *μ*m, the distance between ***E*** cells in the network. For all plots, the data points are plotted as mean +/- standard deviation compiled from 20-100 sweeps.

## S1 Appendix

### Acoustic Space

The general approach for defining acoustic space is as follows. The elements of acoustic space are pure tones of varying frequencies and sound pressures and may be expressed as a product of frequency space 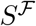 and pressure space 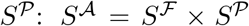. Biologically, the range of audible and discriminable frequencies and sound pressures are limited. To reflect this, the frequency and pressure spaces will each be defined as a topological space where the underlying set is an interval but the topology is coarser than the usual topology.

The frequency space 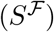 is defined as a half-open interval on the positive real line: 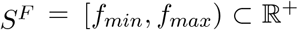, where *f_min_, f_max_* are the minimum and maximum audible frequencies, respectively. To account for the fact that discriminability is limited (i.e. tones of similar frequencies are perceived as the same), 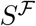 is partitioned into *N_f_* half-open intervals *h_f_* = [*f_α_, f_β_*) = [*f_α_, f_α_* + Δ*f*) where 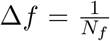 is the length of the interval and loosely represents biologically the minimum frequency difference that can be discriminated. The use of half-open intervals ensures that there is no ‘gap’ between adjacent frequency intervals. The set of all half-open intervals is:

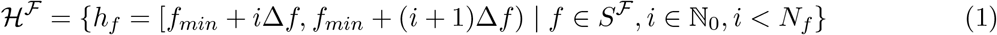

The topology 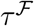 is generated by the elements of 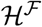:

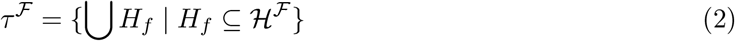

The set of intervals is a partition where each interval is an equivalence class [*f_j_*]: for *f_α_, f_β_* ∈ *S^F^, f_α_* ~ *f_β_* if *f_α_, f_β_* ∈ [*f_i_, f_i_* + Δ*f*). This may be extended so that *f_α_* ~ *f_β_* if *f_α_, f_β_* > *f_max_* and *f_α_* ~ *f_β_* if *f_α_, f_β_* ≤ *f_min_*. Frequencies *f_α_, f_β_* are ‘assigned’ to *f_i_* if they are between *f_i_* and *f_i_* + Δ*f*. Thus, *f_α_, f_β_* are indistinguishable from each other since both are classified as *f_i_*. Each frequency interval is uniquely identified by the minimum point at the closed end. The set of these points is:

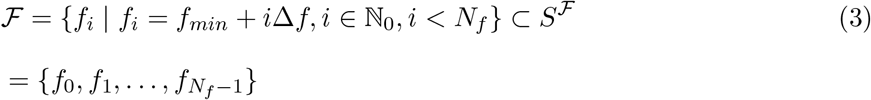

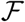 is a discretization of 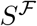 and may be thought of as a set of audible frequencies.

The sound pressure space is constructed similarly to yield:

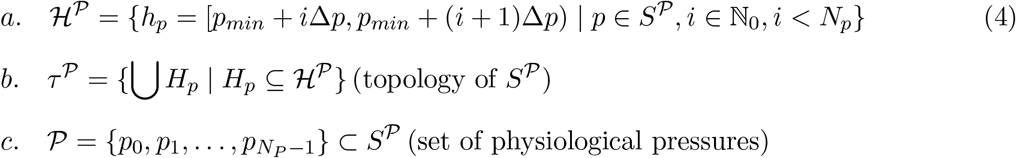

The basis for a topology in 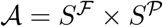 is:

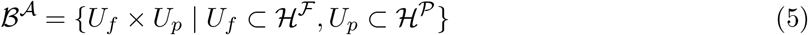

The topology for 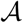 is:

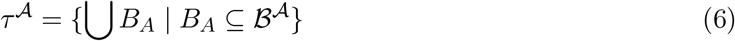

If only the sets of audible frequencies and pressures are considered, then the discretized acoustic space is:

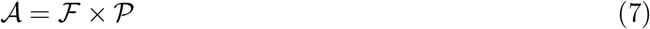

with the discrete topology.

### Neural Space

The neural space 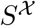 is constructed in a similar fashion as 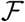 and 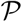, though the topology is slightly different. The idea is to partition 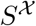 into subintervals that represent neurons and then to combine the subintervals contiguously to form synaptic intervals.

The neural space 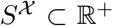 is a half-open interval representing the one dimensional tonotopic axis: 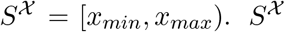 is partitioned into *N_cell_* half-open intervals, each of width Δ*x* and expressed as [*x_α_, x_β_*), *x_β_* > *x_α_* or [*x_α_, x_α_* + Δ*x*). Each interval represents the length (Δ*x*) of a single cell on the tonotopic axis. The set of intervals is expressed as:

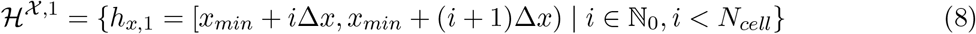

The set 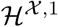 is a partition, which effectively discretizes 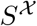 (see section above for a similar treatment). Each interval is uniquely identified by the point at the closed end, which also gives it’s location along the tonotopic axis. The set 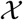 containing these points is:

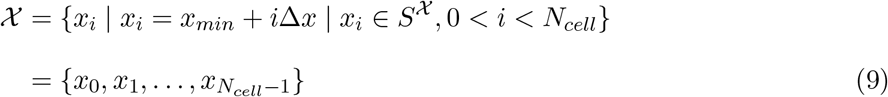

A synaptic interval is also a half-open interval with length λ expressed as:

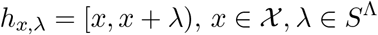

where 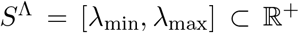 is the range of values that the length can take. The length of the synaptic interval is defined to be a contiguous set of cells so that *h*_*x*,λ_ is an integral multiple of cell length Δ*x*. Letting 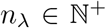 be the number of cells in the interval, the intervals may be re-expressed as:

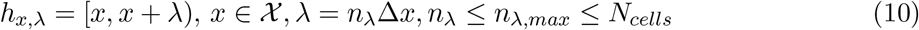

where *n*_λ,*max*_ is the maximum number of cells in the interval. The set containing all possible interval lengths is:

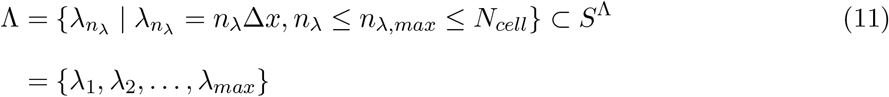

As shown in the next section, sound pressure will be encoded as the length of the intervals. order to ensure a full representation of sound pressure, a ‘buffer’ of *N_cell_* — *n*_λ,max_ cells is need? Without the buffer, the maximum value of last interval is 1 cell. The set of identifying points synaptic intervals is then:

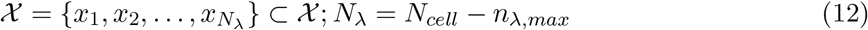

The set of synaptic intervals is expressed as:

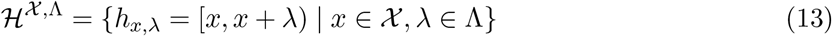

The topology of 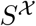 generated by the elements of 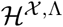 is:

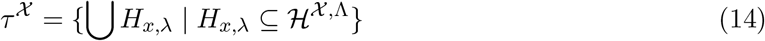

### Mapping from acoustic space to neural space

The aim is to construct a mapping between the acoustic and neural spaces in such a way that allows for both encoding and decoding sounds in neural space.

Note that a pure tone *a_f,p_* corresponds to the ordered pair 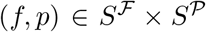 and an interval *h*_*x*, λ_ = [*x,x* + λ) corresponds to the ordered pair 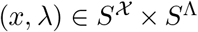. The mapping (*ψ*) from acoustic to neural space is obtained by first defining mappings from frequency space 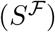 to neural space 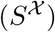 and from pressure space 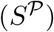 to interval length space (*S*^Λ^):

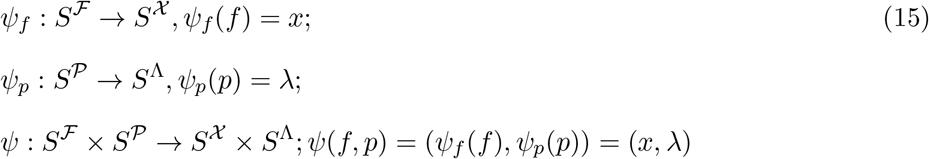

After partitioning, the spaces are effectively discretized into 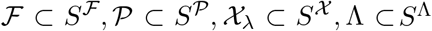. These are the functionally and biological relevant sets as they contain the audible frequencies and pressures that can be mapped into the compartmentalized neural space. Therefore, the mapping can be restricted to these sets:

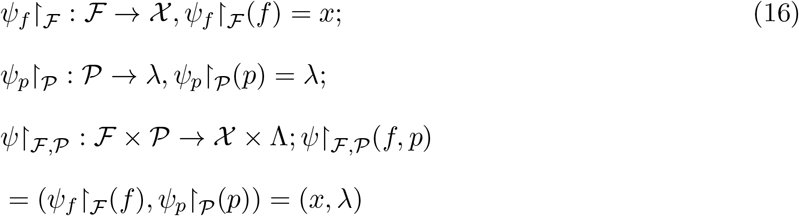

The maps are bijective if 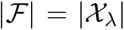 and 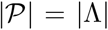 and continuous because the restricted spaces have the discrete topology.

### Algebraic Operations in Neural Space

In neural space, addition (“+”) is also defined as the union. For 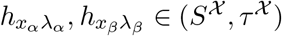

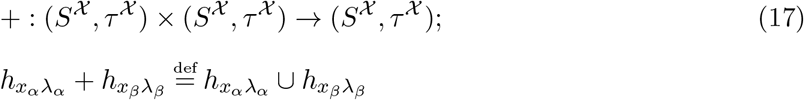

Generally, for any two synaptic intervals 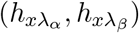, the addition operation yields two possibilities:

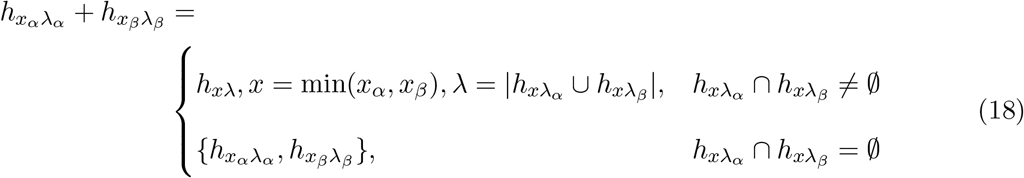

For more than 2 synaptic intervals, the total length λ is given by the Inclusion-Exclusion principle.

The algebraic structure on 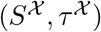 with the addition operation is a monoid since it inherits the algebraic properties of the union operation. In particular, addition is closed because the result is either a longer, half-open interval or two disjoint half-open intervals, either of which is an element of 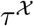. Addition is commutative, associative, has the empty set as the identity element, and does not have an inverse.

Multiplication “·” of synaptic intervals is defined as the set minus operation (\). Given two intervals 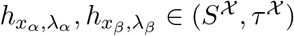 where 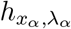 is the multiplier and 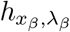 is the multiplicand,

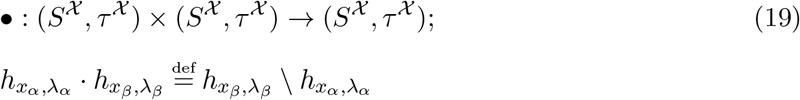

Letting 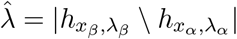, the possible results of multiplication are:

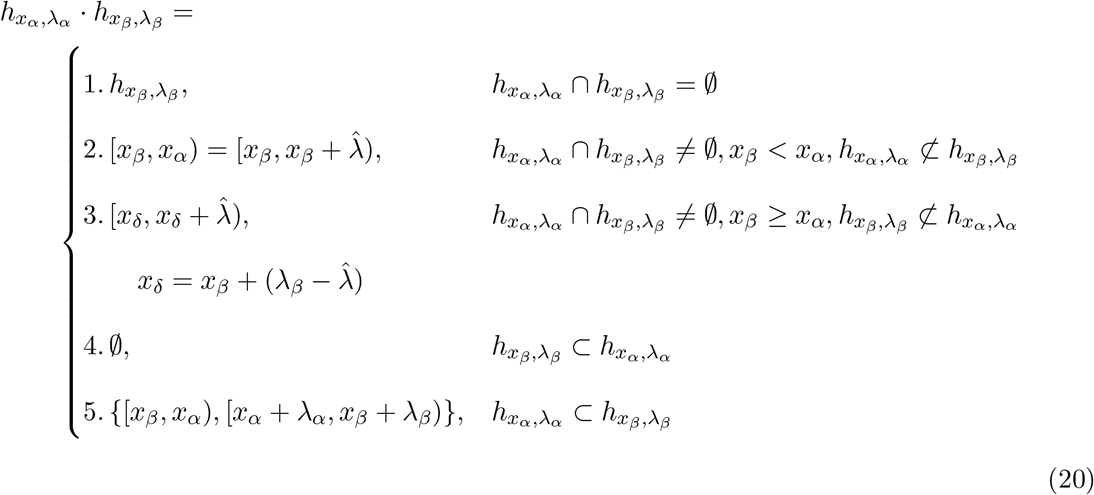

The algebraic structure on 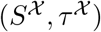 with the multiplication operation is a magma, with properties inherited from the set minus operation. Multiplication is closed because the set minus operation yields an interval that is either the empty set or a half-open interval(s) with a minimum length(s) of Δ*x* and maximum length of λ_*max*_. In either case, the resulting interval is in 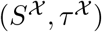. While the 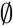 is a left identity element, there is no unique inverse element since the result is the 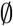 whenever the length of multiplier is greater than or equal to that of the multiplicand. It is also neither commutative nor associative.

#### Proposition 0.0.1.

The *multiplication* operation is left but not right distributive over addition.

*Proof*. **Left distributive**: Let 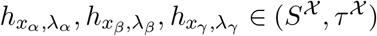

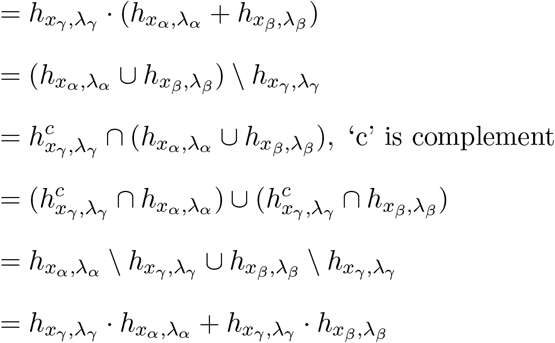

**Not Right distributive**:

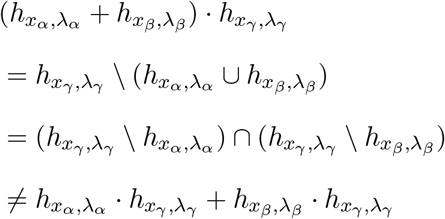

#### Proposition 0.0.2.

Let the representation of a pure tone in neural space be an excitatory interval 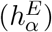 flanked by two inhibitory intervals 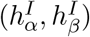 and that of another pure tone be also an excitatory 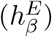 flanked by two inhibitory intervals 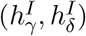, then the combined effect when both tones are introduced simultaneously is not equal to presenting each tone separately and then combining the result. That is, 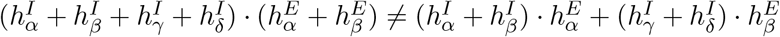.

*Proof*, 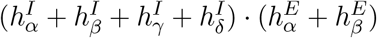 can be re-expressed as:

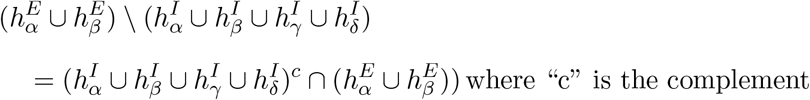

Using a similar argument, 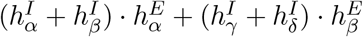 may be re-expressed as:

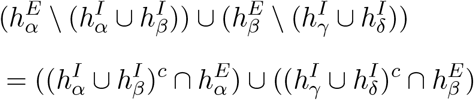

For an element 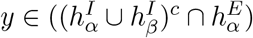, the following statements are true:

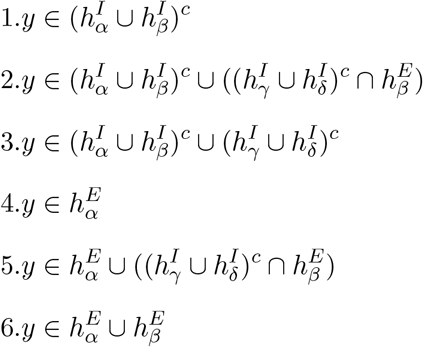

Since 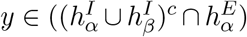, then from lines 3 and 6, 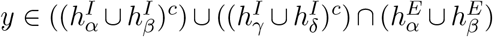.

Therefore,

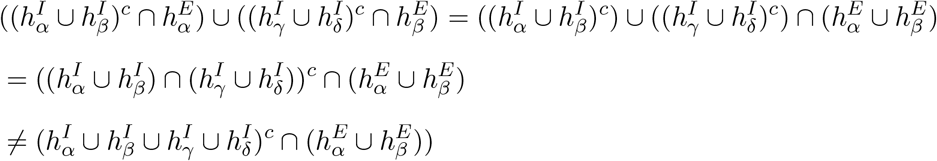

Therefore, 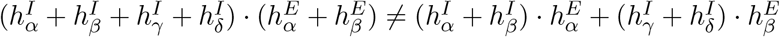

### Comparison with columnar organization

In the mathematical model, the ‘functional’ unit is an interval that can vary in size, depending on the sound intensity. The indistinct borders are consistent with topographical organization of afferents [20]. The expanding border facilitates high resolution representation of both frequency and pressure. Assuming a bijective mapping between acoustic and neural spaces, the maximum number of frequency levels that can be represented is proportional to the number of cells along the tonotopic axis (*N_cell_* and the maximum number of pressure levels is proportional to the maximum number of cells in each interval (*n*_λ,*max*_). The relation between the number of frequency and pressure levels is then given by 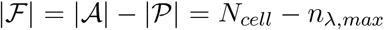 (see eq. 12 in appendix). Hence, the number of frequency levels decreases linearly with the number of pressure levels.

In contrast, consider an alternative neural space that consists of classical columns [18]. The definition of a column has evolved [63] but for illustrative purposes, columns are defined here as having distinct borders; the enclosed neurons have identical receptive fields and hence function as a single unit. In this scheme, the location of the active column encodes frequency and sound pressure is encoded as the number of neurons that become active during the stimulus (assuming a brief stimulus so that neurons are effectively binary).

Construction of the neural space is as above except that the flexible synaptic intervals are replaced with half-open intervals, termed columns, spanning a *fixed* number of cells *n_col_*. The number of columns that fit into the neural space with *N_cell_* neurons is therefore 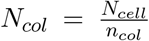. Assuming a bijective mapping from frequency space to neural space, the number of frequency levels 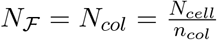. Because the number of cells in a column (*n_col_*) is fixed, pressure will be encoded as the number of cells that become active within a column during a stimulus. Assuming a bijective map, the number of pressure levels is 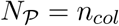.

Hence, the number of frequency levels is inversely proportional to the number of pressure levels: 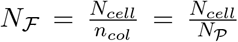. Compared with the model with flexible synaptic intervals, the number of frequency levels drops off much faster with the number of pressure levels.

### Experimental evidence

**Supporting Fig. S1:**
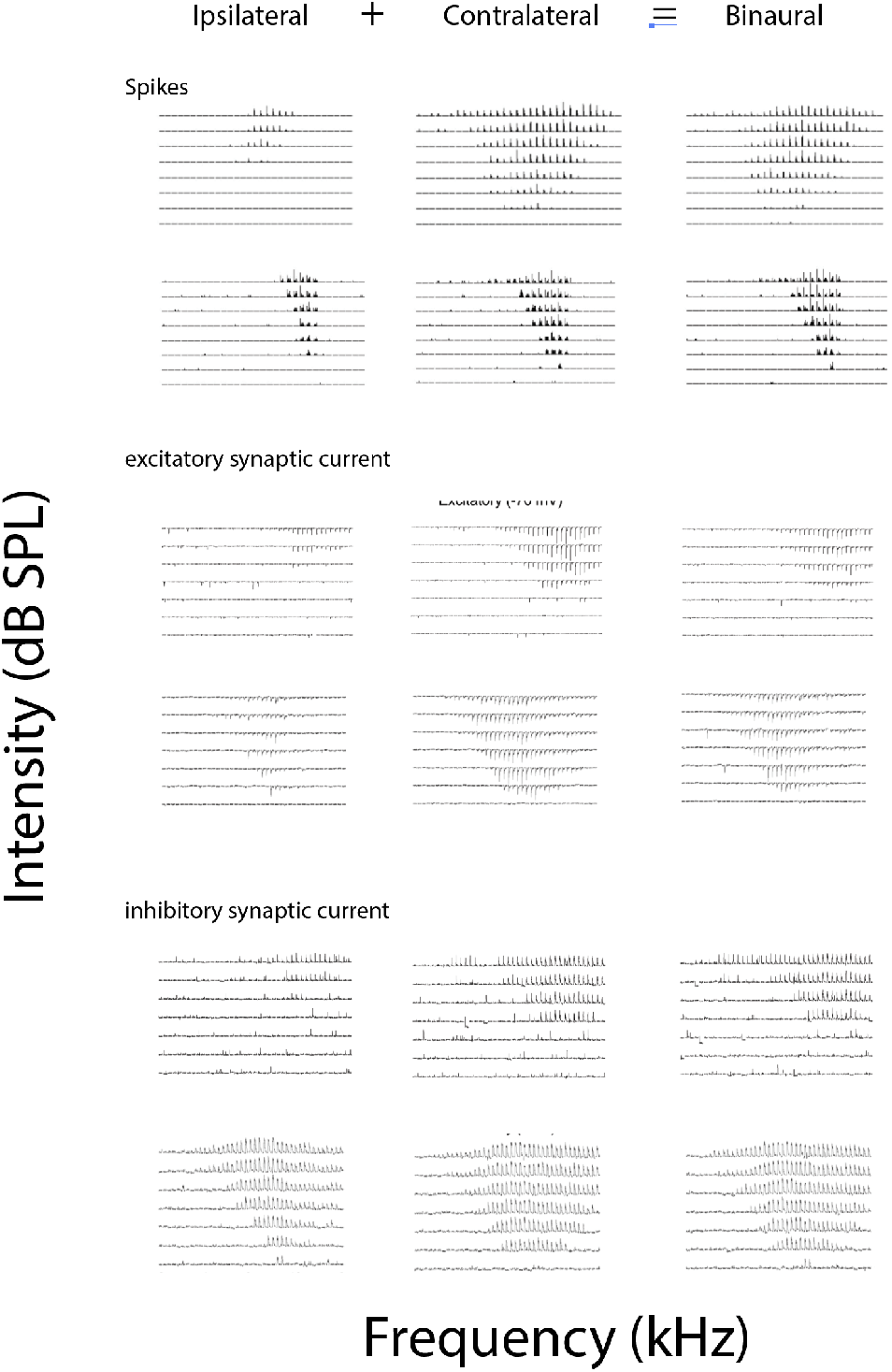
Experimental evidence for Addition operation. Data adapted from reference [33]. Extracellular (top) and whole-cell voltage clamp (middle, bottom) recording from the central nucleus of the Inferior colliculus, which receives binaural inputs. The frequency response areas (FRA) were constructed by presenting pure tones of varying frequencies (abscissas) and intensities (ordinates) and documenting the responses. Tones were delivered ipsilaterally (left column), contralaterally (middle), and binaurally (right). The FRAs can be taken as an indicator of the spatial extent of synaptic field and activated area in the following manner. Note that the FRAs are “V-shaped”. At low intensities, neurons fire only when the tone frequencies are near its preferred frequency (tip of the V). At higher intensities, the range of frequencies that evoke firing increases substantially. If adjacent neurons have comparably-shaped FRAs but have slightly different preferred frequencies, an increase in intensity would translate to an increase in the spatial extent of activated neurons. The model predicts that because the contralateral response is greater than the ipsilateral response, the response evoked with binaural stimulation should equal that of contralateral stimulation. In all cases, the binaural FRAs was nearly identical to the contralateral FRAs. *Permission pending*.

### Algebra of loudness summation

A multi-frequency stimulus is described as a set of frequencies 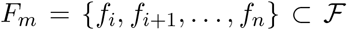. In neural space, the stimulus generates a set of excitatory intervals *H_m_* that is the union of the individual intervals generated by each tonal component: 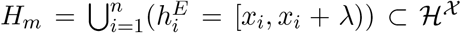, where λ is the length of each interval.

#### Definition 0.0.1.

A tone with frequency *f_d_* in multi-frequency stimulus is ‘dominant’ if, in addition to an excitatory interval 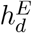, it generates an inhibitory interval *h^I^* that is larger than those generated by others in neural space.

Physiologically, the dominant frequency *f_d_* could be the center frequency that identifies a CB [42] or the lowest frequency tone of a complex stimulus [30]. It is assumed that every multifrequency stimulus has a dominant frequency, which generates the synaptic interval 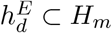.

#### Definition 0.0.2.

The inhibitory interval is in the lateral inhibitory configuration if 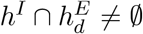 and there exists a half-open interval 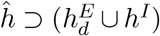 so that if 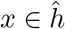, then 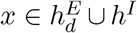. This ensures that the excitatory and inhibitory intervals abut each other with no space in between. Note that this definition applies whether *h^I^* is to one side of 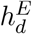 or is split into two that occupies either side (as in the main text) 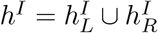.

The interaction of 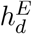 and *H_m_* described by the algebraic operation: *h_l_* = *h^I^* · *H_m_*. The length |*h_l_*| is taken to be the proxy for loudness perception.

#### Proposition 0.0.3.

In a lateral inhibition configuration, increasing the size of *H_m_* does not increase *h_l_* as long as *H_m_* remains within (is a subset of) a ‘critical interval’ *h_CI_*.

*Proof. H_m_* can be divided in 3 disjoint sets according to their relation to 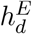 and *h^I^*.

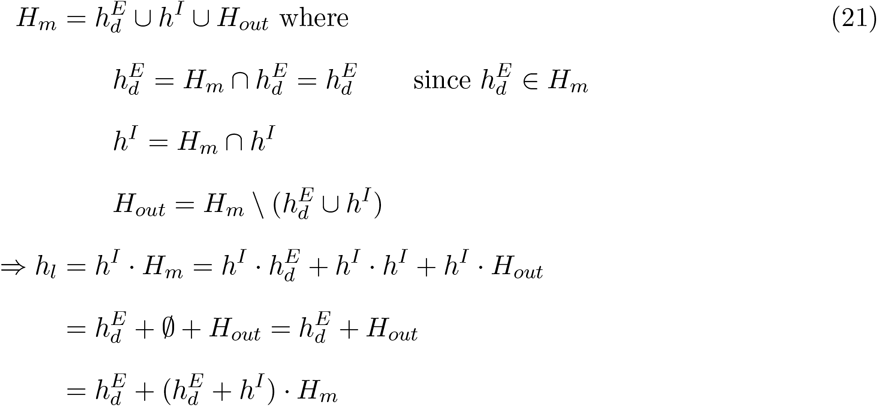

Thus, *h_l_* is equal to 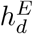 as long as 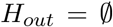 or when 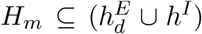. The critical interval is therefore 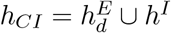.

#### Definition 0.0.3.

The critical band (CB) is difference between the highest frequency and lowest frequency in *F_m_* that satisfies *H_m_* ⊆ *h_CI_*.

#### Proposition 0.0.4.

The critical band does not change with stimulus intensity. In the following, an increase in stimulus intensity is represented by an increase in the length of each excitatory interval by an amount Δ*h*.

*Proof*. Let 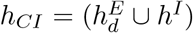, which is a single interval 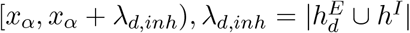 if in the lateral inhibition configuration. As per above, 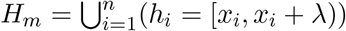. For 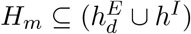 to be true, the starting point of the first interval *x*_1_ ≥ *x_α_* and the starting point *x_n_* of the last point must be such that *x_n_* + λ ≤ *x_α_* + λ_*d,inh*_. In frequency space, *x*_1_ is associated with a frequency *f*_1_ and *x_n_* with *f_n_* so the CB is *f_n_* – *f*_1_.

Increasing the lengths all of the excitatory intervals by Δ*h* effectively increases 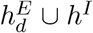 also by Δ*h*. For example, the union of two successive intervals *h*_1_ = [*x*_1_, *x*_1_ + λ), *h*_2_ = [*x*_2_, *x*_2_ + λ) is [*x*_1_, *x*_1_ + λ) ∪ [*x*_2_, *x*_2_ + λ) = [*x*_1_, *x*_2_ + λ). Increasing each interval by Δ*h* results in [*x*_1_, *x*_1_ + λ + Δ*h*) ∪ [*x*_2_, *x*_2_ + λ + Δ*h*) = [*x*_1_, *x*_2_ + λ + Δ*h*). Therefore, the condition for starting point of the first interval is still *x*_1_ ≥ *x_α_* and that for the starting point of the last interval is *x_n_* + λ + Δ*h* ≤ *x_α_* + λ_*d,inh*_ + Δ*h*. Hence, *n* does not change. If *H_m_* is due to a set of frequencies *F_m_* = {*f*_1_, *f*_2_,…, *f_n_*}, then then CBis still *f_n_* – *f*_1_.

The total length (*h_l_*) resulting from the operations are shown graphically for band-limited white noise with increasing bandwidth (main text) and for a complex stimulus with 4 tones with increasing spacing between frequencies (Supporting Fig. S2)

**Supporting Figure S2:**
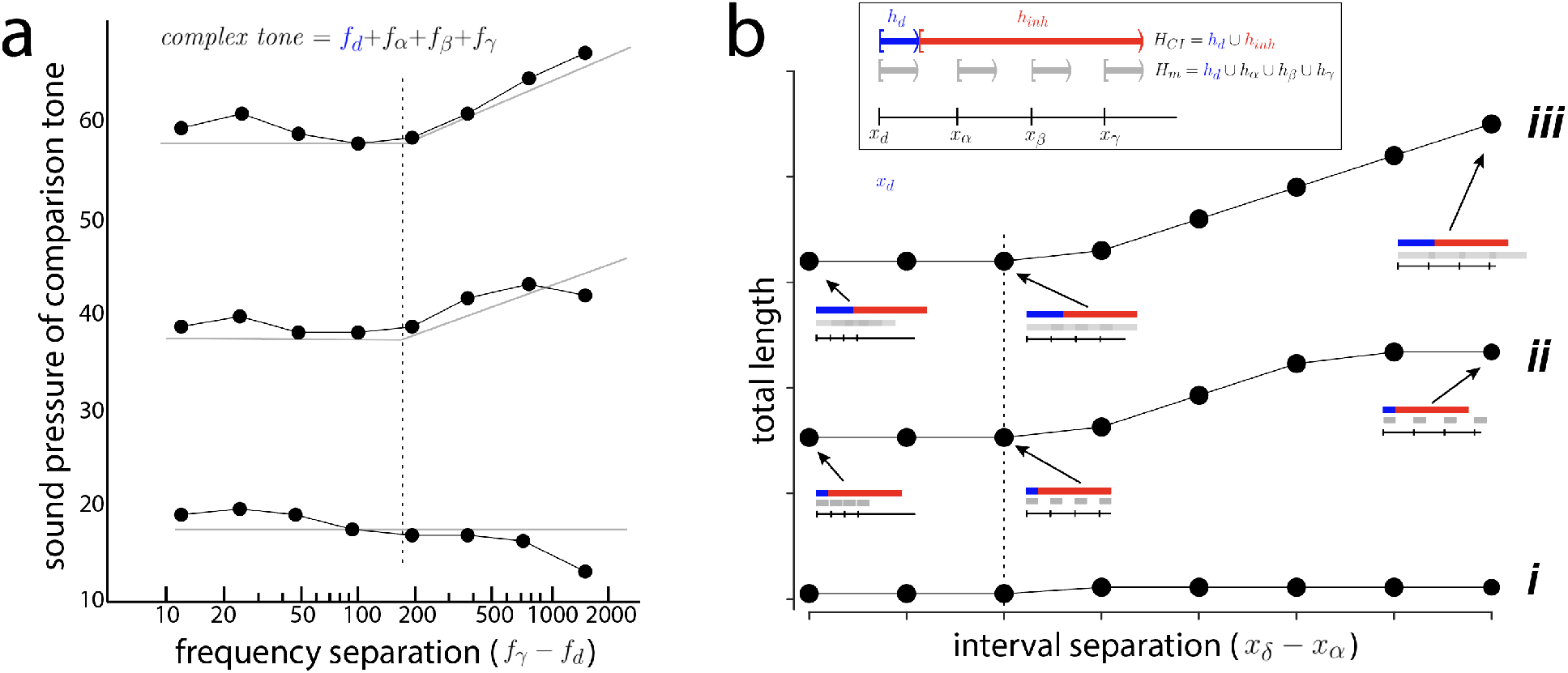
Algebra of loudness summation. ***a***, Perceived loudness when subjects were presented with a complex tone with 4 frequency components spaced evenly apart at 3 sound pressure levels. Subjects were asked to increase the intensity of a single tone (ordinate) until it was as loud as the complex tone. The frequency separation was systematically increased (abscissa). Dotted vertical line corresponds to critical band. Figure adapted from [44]. ***b***, Predicted interval lengths resulting from the interaction of 4 tones delivered simultaneously (inset). ***Boxed inset***, The tones generate 4 excitatory intervals (*H_m_*, gray) one of which is dominant (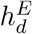, *blue*). The dominant tone also generates an inhibitory synaptic interval (red). Plot shows resultant length (*h_l_*) after the operations as the frequency spacing between intervals in *H_m_* is increased (abscissa). Dotted vertical line marks deviation from constant value.

